# Acetylcholine is released in the basolateral amygdala in response to predictors of reward and enhances learning of cue-reward contingency

**DOI:** 10.1101/2020.04.14.041152

**Authors:** Richard B. Crouse, Kristen Kim, Hannah M. Batchelor, Rufina Kamaletdinova, Justin Chan, Prithviraj Rajebhosale, Steven T. Pittenger, Lorna W. Role, David A. Talmage, Miao Jing, Yulong Li, Xiao-Bing Gao, Yann S. Mineur, Marina R. Picciotto

## Abstract

The basolateral amygdala (BLA) is critical for associating initially neutral cues with appetitive and aversive stimuli and receives dense neuromodulatory acetylcholine (ACh) projections. We measured BLA ACh signaling and principal neuron activity in mice during cue-reward learning using a fluorescent ACh sensor and calcium indicators. We found that ACh levels and activity of nucleus basalis of Meynert (NBM) cholinergic terminals in the BLA (NBM-BLA) increased sharply in response to reward-related events and shifted as mice learned the tone-reward contingency. BLA principal neuron activity followed reward retrieval and moved to the reward-predictive tone after task acquisition. Optical stimulation of cholinergic NBM-BLA terminal fibers during cue-reward learning led to more rapid learning of the cue-reward contingency. These results indicate that BLA ACh signaling carries important information about salient events in cue-reward learning and provides a framework for understanding how ACh signaling contributes to shaping BLA responses to emotional stimuli.

## Introduction

Learning how environmental stimuli predict the availability of food and other natural rewards is critical for survival. The basolateral amygdala (BLA) is a brain area necessary for associating cues with both positive and negative valence outcomes (Baxter & Murray, 2002; Janak & Tye, 2015; LeDoux et al., 1990). Recent work has shown that genetically distinct subsets of BLA principal neurons encode the appetitive and aversive value of stimuli (J. Kim et al., 2016). This encoding involves the interplay between principal neurons, interneurons, and incoming terminal fibers, all of which need to be tightly regulated to function efficiently.

The neuromodulator acetylcholine (ACh) is released throughout the brain and can control neuronal activity via a wide range of mechanisms. ACh signals through two families of receptors (nicotinic, nAChRs and muscarinic, mAChRs) that are differentially expressed on BLA neurons as well as their afferents (Picciotto et al., 2012). ACh signals through these receptors to increase signal-to-noise ratios and modify synaptic transmission and plasticity in circuits involved in learning new contingencies (Picciotto et al., 2012), especially in areas that receive dense cholinergic input, like the BLA (Woolf, 1991; Zaborszky et al., 2012).

The basal forebrain complex is a primary source of ACh input to the BLA. In particular, the nucleus basalis of Meynert (NBM) sends dense cholinergic projections to the BLA (Woolf, 1991; Zaborszky et al., 2012). Optical stimulation of BLA-projecting cholinergic terminal fibers (NBM-BLA) during fear conditioning is sufficient to strengthen fear memories (Jiang et al., 2016) and may support appetitive behavior (Aitta-aho et al., 2018). Cholinergic NBM neurons increase their firing in response to both rewarding and aversive unconditioned stimuli (Hangya et al., 2015). A recent study has also demonstrated that NBM cells fire in response to a conditioned stimulus during trace fear conditioning, indicating that ACh signaling may be involved in learning about cues that predict salient outcomes (Guo et al., 2019).

We hypothesized that ACh signaling in the BLA is a critical neuromodulatory signal that responds to both unconditioned stimuli and cues that gain salience, thereby coordinating activity in circuits necessary for learning cue-reward contingencies. To test this hypothesis, we measured relative levels of BLA ACh (ACh signaling), cholinergic NBM-BLA terminal fiber activity (BLA ACh signal origin), and the activity of BLA principal neurons (BLA output) across all phases of learning in an appetitive operant learning task to evaluate how BLA output and ACh signaling are related to behavioral performance in this paradigm. We then optically stimulated cholinergic NBM fibers locally in the BLA while mice learned to nose poke in response to an auditory cue to receive a food reward to determine if accelerating the increase in ACh signaling that occurs as mice learn the task would enhance performance. We also pharmacologically blocked different ACh receptors during the learning task to determine the subtypes involved, and varied the timing of optical stimulation of cholinergic NBM-BLA terminal fibers to determine whether time-locked ACh release with the reward-predictive cue is necessary for the improvement of the task performance. These studies provide a novel framework for understanding how NBM ACh signaling in the BLA is recruited during perception of novel stimuli and how it contributes to linking previously neutral cues to predictions about future salient outcomes.

## Results

### Acetylcholine release in the BLA occurs at salient points in the cue-reward learning task and shifts as mice learn the cue-reward contingency

The BLA is critical for learning that previously neutral cues can predict future punishments or rewards and for assigning valence to those cues (Baxter & Murray, 2002; Janak & Tye, 2015). The BLA receives dense cholinergic input (Woolf, 1991; Zaborszky et al., 2012) and we speculated that, since ACh signaling is involved in both attention and several types of learning (Picciotto et al., 2012), it could be essential for learning about cues that predict salient events, such as reward delivery. Based on data showing that ACh neurons fire in response to unexpected or salient events (Hangya et al., 2015), we also hypothesized that ACh release might vary as mice learn a cue-reward contingency. Therefore, we designed a cue-reward learning task in which food-restricted mice were trained to perform a nose poke when signaled by a cue (tone) to receive a palatable reward (Ensure) on a 30 sec variable intertrial interval (ITI) (**Fig. 1A-D**). We injected adeno-associated virus (AAV) carrying an improved version of the fluorescent ACh sensor GRAB_ACh3.0_ (ACh3.0; (Jing et al., 2018, 2019) construct into the BLA of mice and implanted an optical fiber above the BLA to record ACh signaling during the cue-reward learning task (**Fig. 2A + S2.1A**).

**Fig. 1.**
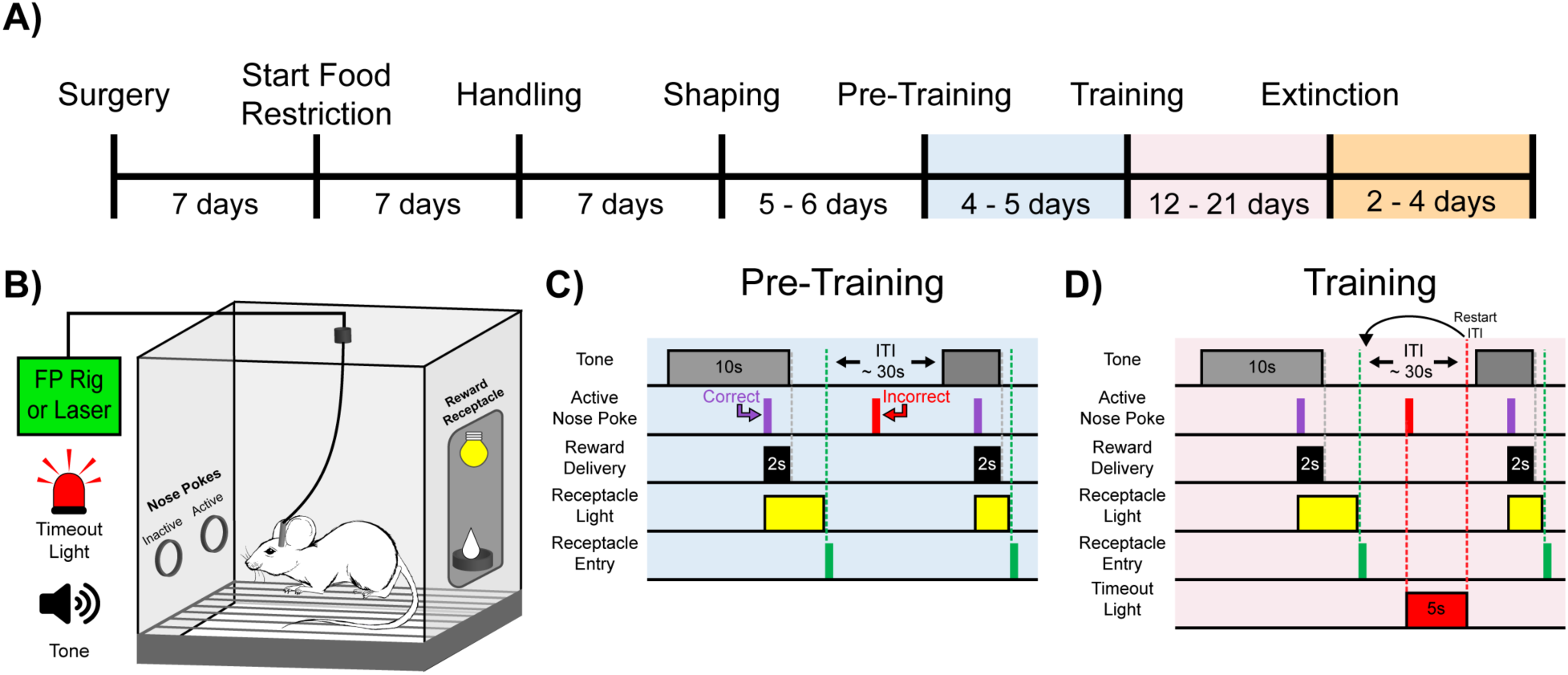
Experimental Timeline and Cue-Reward Learning Paradigm. A) <u>Experimental timeline.</u> Mice began food restriction 7 days after surgery and were maintained at 85% free-feeding body weight for the duration of the experiment. After 7 days of handling, 5-6 days of behavioral shaping prepared the mice for the cue-reward learning task (Pre-Training through Extinction). B) <u>Behavioral chamber setup.</u> Mice were placed in modular test chambers that included two nose poke ports on the left wall (Active and Inactive) and the Reward Receptacle on the right wall. A tone generator and timeout light were placed outside the modular test chamber. For fiber photometry (FP) and optical stimulation (Laser) experiments, mice were tethered to a patch cord(s). <u>C-D) Details of the Cue-Reward Learning Paradigm</u> C) In Pre-Training, an auditory tone was presented on a variable interval 30 schedule (VI30), during which an active nose poke yielded Ensure reward delivery but there was no consequence for incorrect nose pokes (active nose pokes not during tone). D) Training was identical to Pre-Training, except incorrect nose pokes resulted in a 5 sec timeout, signaled by timeout light illumination, followed by a restarting of the intertrial interval (ITI).

**Fig. 2.**
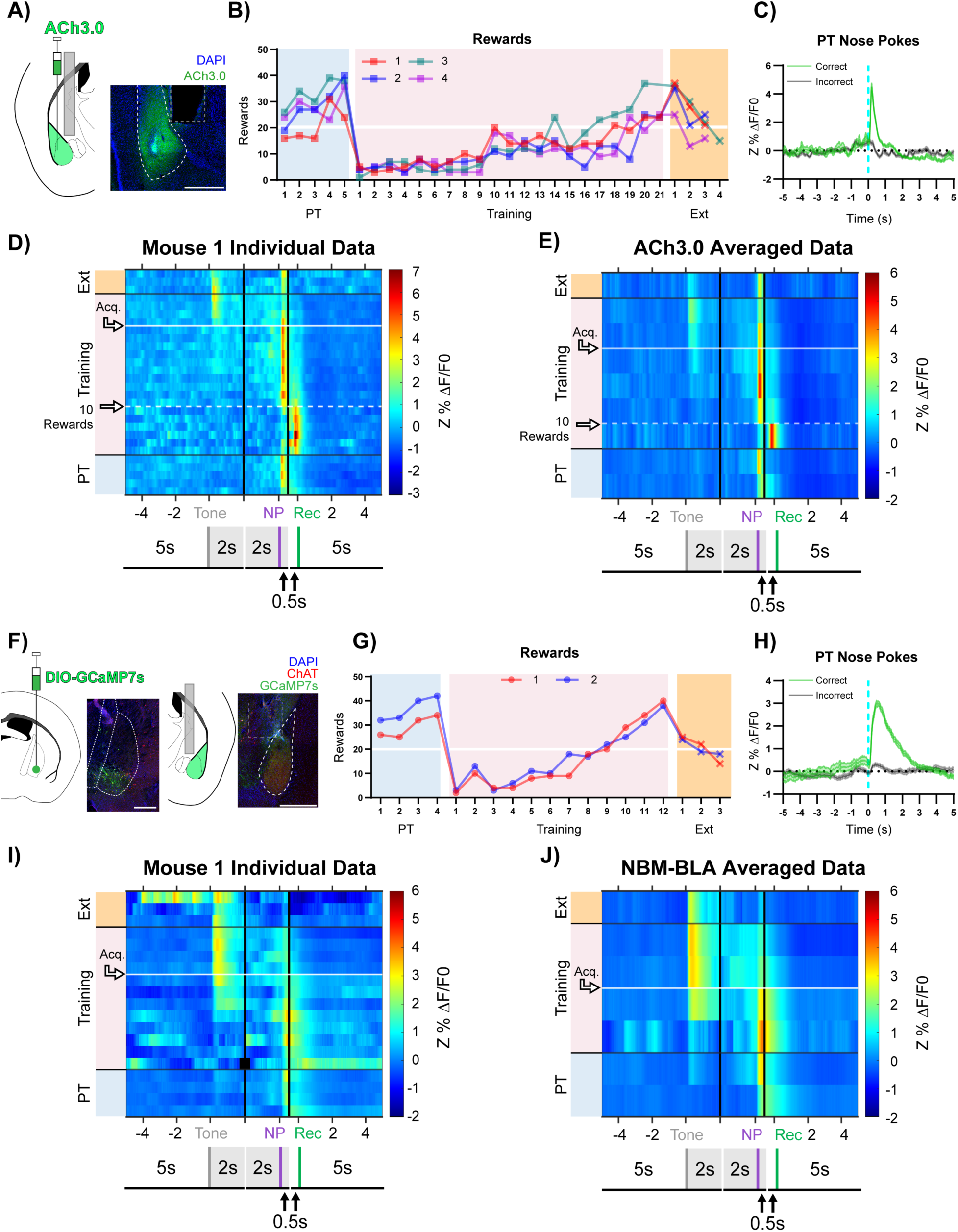
Basolateral Amygdala (BLA) ACh Signaling Aligns with Salient Events During Reward Learning A) <u>Diagram and example of injection and fiber placement sites in the BLA for recording from mice expressing a fluorescent acetylcholine sensor (ACh3.0)</u>. Left: Diagram of BLA ACh3.0 injection and fiber tip placement. Right: Representative coronal brain slice with fiber tip and ACh3.0 expression. Blue: DAPI, Green: ACh3.0. White dashed line: BLA outline. Grey dashed rectangle: fiber track. Scale = 500 µm. Individual fiber placements are shown in **Fig. S2.1A**. B) <u>Behavioral responding of mice expressing ACh3.0 in BLA</u>. Individual mice acquired the task at different rates as measured by rewards earned. Horizontal white line: acquisition threshold, when a mouse began to earn ∼20 rewards consistently in Training. Incorrect nose pokes 1shown in **Fig. S2.2A**. Pre-Training (PT): blue shaded area, Training: pink shaded area, Extinction (Ext): orange shaded area. C) <u>Fluorescence traces from BLA of ACh3.0-expressing mice</u>. A substantial increase in fluorescence representing BLA ACh release coincided with correct (green line) but not incorrect (grey line) nose pokes on last day of PT (data are shown from Mouse 1). Mean ± SEM, correct (n = 24), incorrect (n = 58). Traces of signal and reference channels (%ΔF/F0) during nose pokes are shown in **Fig. S2.1B-C**. Incorrect nose pokes on last day of PT vs Training Day 1 shown in **Fig. S2.2B.** D) <u>Heatmap of BLA ACh signaling in mouse 1 across all training phases, aligned to tone onset (Tone), correct nose poke (NP), and receptacle entry (Rec)</u>. Each row is the average of rewarded trials across a training session. White dashed horizontal line: first Training day earning 10 rewards. Horizontal white line: acquisition threshold, when a mouse began to earn ∼20 rewards consistently in Training. Black horizontal lines: divisions between training phases. Black vertical lines: divisions between breaks in time to allow for variable latencies in tone onset, correct nose poke, and receptacle entry (reward retrieval). Individual data for mice 2-4 in **Fig. S2.1D-F**. Incorrect nose pokes heatmaps for individual mice shown in **Fig S2.2C-F.** E) <u>Heatmap of BLA ACh signaling averaged across mice</u>. Signal aligned as in D) with a selection of data from key days in the behavioral paradigm shown. From bottom to top: PT Day 1, PT Day 5, Training Day 3, First Training day earning 10 rewards (white dashed horizontal line), Training Day 13, Training Day 15, Acquisition day (white horizontal line), Last Training Day, Last Extinction Day. Black horizontal lines: divisions between training phases. Black vertical lines: divisions between breaks in time to allow for variable latencies in tone onset, correct nose poke, and receptacle entry. Incorrect nose poke heatmaps averaged across mice shown in **Fig. S2.2G** F) <u>Diagram and example of Nucleus Basalis of Mynert (NBM)-BLA terminal fiber recordings</u>. Left: DIO-GCaMP7s was injected in the NBM of ChAT-IRES-Cre mice, individual injection sites are shown in **Fig. S2.3A**. Representative coronal brain slice showing GCaMP7s expression. White dashed lines: internal capsule and globus pallidus outlines. Blue: DAPI, Green: GCaMP7s, Red: ChAT. Scale = 500 µm; separate channels shown in **Fig. S2.3C**. Right: An optical fiber was implanted above the ipsilateral BLA, individual fiber placements are shown in **Fig. S2.3B**. Representative coronal brain slice showing GCaMP7 expression and fiber tip placement. White dashed line: BLA outline. Grey dashed rectangle: fiber tract. Blue: DAPI, Green: GCaMP7s, Red: ChAT. Scale = 500 µm; separate channels shown in **Fig. S2.3D**. G) <u>Behavioral responding of mice during NBM-BLA terminal fiber recordings.</u> Individual mice acquired the task at different rates as measured by rewards earned. White horizontal line: acquisition threshold, when a mouse began to earn ∼20 rewards consistently in Training. Incorrect nose pokes shown in **Fig. S2.4A.** H) <u>NBM-BLA terminal fiber activity is similar to ACh3.0 recordings</u>. NBM-BLA terminal fiber activity coincided with correct (green line) but not incorrect (grey line) nose pokes on last day of PT (data shown for Mouse 1). Mean ± SEM, correct (n = 42), incorrect (n = 101). Signal and reference channels (%ΔF/F0) during nose pokes are shown in **Fig. S2.3E-F**. Incorrect nose pokes on last day of PT vs Training Day 1 shown in **Fig. S2.4B.** See **Fig. S2.5A-H** for ACh3.0 and NBM-BLA terminal fiber recordings in the same mouse. I) <u>Heatmap of NBM-BLA terminal fiber activity in mouse 1 across all training phases, aligned to tone onset (Tone), correct nose poke (NP), and receptacle entry (Rec)</u>. Each row is the average of rewarded trials across a training session. Horizontal white line: acquisition threshold, when a mouse began to earn ∼20 rewards consistently in Training. Black horizontal lines: divisions between training phases. Black vertical lines: divisions between breaks in time to allow for variable latencies in tone onset, correct nose poke, and receptacle entry (reward retrieval). Blanks in the heatmaps indicate time bins added for alignment. Mouse 2 individual data shown in **Fig. S2.3G**. Incorrect nose pokes heatmaps for individual mice shown in **Fig S2.4C-D.** J) <u>Heatmap of NBM-BLA terminal fiber activity averaged across mice</u>. Signal aligned as in D-E) with a selection of key days shown, from bottom to top: PT Day 1, PT Day 4, Training Day 3, Training Day 6, Acquisition day (white horizontal line), Last Training Day, Last Extinction Day. Black horizontal lines: divisions between training phases. Black vertical lines: divisions between breaks in time to allow for variable latencies in tone onset, correct nose poke, and receptacle entry (reward retrieval). Incorrect nose poke heatmaps averaged across mice shown in Fig. S2.4E.

During the Pre-Training phase of the task, mice received reward and cue light presentation for performing a nose poke in the active port during tone presentation (**Fig. 1C**, purple active nose poke coincident with tone) but there was no consequence for an incorrect nose poke (**Fig. 1C**, red active nose poke not coincident with tone). Animals quickly learned to make a high number of responses over the course of each Pre-Training session. In this paradigm, mice obtained most available rewards by day 5 of Pre-Training (**Fig. 2B**, blue shaded region). However, this phase of training did not promote learning of the cue-reward contingency, (i.e. that they should only nose poke during tone presentation) seen by the high number of incorrect nose pokes (**Fig. S2.2A**, blue shaded region). Mice performed roughly 8-fold more incorrect nose pokes than correct nose pokes, suggesting that mice were not attending to the task contingency. The Training phase of the task was identical to Pre-Training except incorrect nose pokes resulted in a 5 sec timeout, during which the house light was illuminated, that concluded with a restarting of the ITI timer (**Fig. 1D**, red active nose poke not coincident with tone). On day 1 of the Training phase, all animals earned fewer rewards (**Fig. 2B**, pink shading) and, while still high, incorrect nose pokes dropped (**Fig. S2.2A**, pink shading). Animals that did not progress to the cut off for acquisition by day 9 (defined as consistently earning 20 or more rewards per session, **Fig. 2B**, white horizontal line) were moved to a 20 sec variable ITI to promote responding (**Fig. 2B**, pink shading day 10). Following the change in ITI, mice acquired the cue-reward behavior at different rates. After acquisition, animals were switched to Extinction training in which correct nose pokes did not result in reward delivery, and all mice decreased nose poke responding (**Fig. 2B** + **Fig. S2.2A**, orange shading).

During Pre-Training, when there were high numbers of both correct and incorrect nose pokes, there was a large increase in ACh release following correct nose pokes, which were followed by reward delivery and cue light, but not incorrect nose pokes (**Fig. 2C + Fig. S2.1 B-C**). ACh release occurred in response to different events as mice learned the task (example data for each mouse is shown in **Fig. 2D** + **Fig. S2.1D-F** and averaged data across all mice at key time points in the task is shown in **Fig. 2E**). During Pre-Training rewarded trials, the highest levels of ACh release occurred immediately after correct nose pokes (NP), with a smaller peak at the time of reward retrieval (entry into the reward receptacle, Rec). As Training began, the ACh release during reward trials shifted dramatically toward the time of reward retrieval, likely because the animals were learning that many nose poke events did not result in reward delivery. Incorrect nose pokes that triggered a timeout were also followed by a modest increase in BLA ACh levels (**Fig. S2.2B-G**). As mice began to learn the contingency (**Fig. 2E**, 10 rewards, white horizontal dashed line), the peak ACh release during rewarded trials shifted back to the time of the correct nose poke response but the peak following incorrect nose pokes remained (**Fig. S2.2C-G**). As animals approached the acquisition criterion (**Fig. 2E**, Acq., white horizontal line), ACh level also increased at the time of the tone and decreased at the time of reward, suggesting that as animals learned the cue-reward contingency, the tone became a more salient event. After task acquisition, the increase in ACh following correct nose pokes remained but was diminished, while incorrect nose pokes no longer elicited apparent ACh release. During Extinction, ACh release to tone onset diminished.

In order to determine the source of the ACh released in the BLA during cue-reward learning, we recorded calcium dynamics as a measure of cell activity of ChAT^+^ NBM terminal fibers in the BLA (NBM-BLA), since the NBM is a major source of cholinergic input to the BLA (Jiang et al., 2016; Woolf, 1991; Zaborszky et al., 2012). We injected AAV carrying a Cre-recombinase-dependent, genetically-encoded calcium indicator (DIO-GCaMP7s) into the NBM of ChAT-IRES-Cre mice and implanted an optical fiber above the ipsilateral BLA (**Fig. 2F + Fig. S2.3A-D)**. Mice in this cohort learned in a similar fashion (**Fig. 2G + Fig. S2.4A**) but met the acquisition criteria faster than mice in the ACh3.0 sensor recording experiment because aspects of the behavioral setup were optimized for the imaging apparatus. As with the recording of ACh3.0 sensor, there was a dramatic difference in NBM-BLA cholinergic terminal activity between correct vs incorrect nose pokes (**Fig. 2H + Fig. S2.3 E-F**). NBM-BLA cholinergic terminal activity evolved across phases of the reward learning task as was seen for ACh levels in the BLA (data for each mouse shown in **Fig. 2I + S2.3G**, averaged across all mice at key time points in the task shown in **Fig. 2J**). Strikingly, NBM-BLA cholinergic terminal activity followed correct nose pokes in Pre-Training and shifted primarily to tone onset as mice learned the contingency during Training. Incorrect nose pokes that resulted in a timeout in Training sessions were followed by a modest increase in NBM-BLA cholinergic terminal activity before task acquisition, similar to what was seen for ACh levels (**Fig. S2.4 B-E**). During Extinction, activity of NBM-BLA terminals following tone onset diminished.

In order to record NBM-BLA cholinergic terminal activity and BLA ACh levels simultaneously in the same mouse, we injected AAV carrying a construct for Cre-recombinase dependent red-shifted genetically-encoded calcium indicator (DIO-jRCaMP1b) into the NBM of ChAT-IRES-Cre mice, ACh3.0 sensor into the ipsilateral BLA, and implanted a fiber above the BLA (**Fig. S2.5A-E**, mouse 1). DIO-jRCaMP1b was also injected into the NBM of a wild type littermate so Cre-mediated recombination would not occur to control for any crosstalk between the ACh3.0 and jRCaMP1b channels. We found that NBM-BLA cholinergic terminal activity coincided with ACh levels (**Fig. S2.5F-G**). Importantly, this relationship between ACh release and NBM-BLA terminal fiber activity was not explained by signal crosstalk (**Fig. S2.5H**), further indicating that the BLA ACh measured comes at least in part from the NBM.

### BLA principal neurons respond to reward availability and follows cue-reward learning

Glutamatergic principal cells are the primary output neurons of the BLA (Janak & Tye, 2015), and their firing is modulated by NBM-BLA cholinergic signaling (Jiang et al., 2016; Unal et al., 2015). BLA principal neurons can increase their firing in response to cues as animals learn cue-reward contingencies (Sanghera et al., 1979; Schoenbaum et al., 1998; Tye & Janak, 2007). To determine whether ACh modulates principal neuron activity during cue-reward learning, we injected AAV carrying a Cre-recombinase dependent genetically encoded calcium indicator (DIO-GCaMP6s) into the BLA of CaMKIIα-Cre mice to record BLA principal cell activity during the learning task (**Fig. 3A + S3.1A**). As was seen for BLA ACh levels, there was a substantial difference in BLA principal cell activity following correct and incorrect nose pokes on the last day of Pre-Training (**Fig. 3B**). However, the activity peaked later after the nose poke response (∼2.5 sec) compared to the ACh3.0 signal (∼0.5 sec) and appeared to align more tightly with reward retrieval (**Fig. S3.1B**). As mice learned the task (**Fig. 3C + Fig. S3.2A**), BLA principal cell activity increased first in response to reward and after acquisition of the task, to the reward-predictive cue (individual data for each mouse shown in **Fig. 3D** + **Fig. S3.1E-F,** and averaged data across all mice at key time points in task is shown in **Fig. 3E**). During Pre-Training, the highest levels of BLA principal cell activity followed reward retrieval. In addition, during the first few days of Training, BLA principal cell activity after reward retrieval was higher than it was during Pre-Training, and the magnitude of response decreased as mice learned the contingency and earned more rewards, ultimately reaching similar intensity to that observed during Pre-Training. Concurrently, as mice approached acquisition of the task (**Fig. 3C**, white horizontal line), BLA principal cell activity increased in response to tone onset (**Fig. 3D-E + Fig. S3.1E-F**, Acq., white horizontal line), suggesting that the recruitment of BLA principal cell activity likely reflects the association of the cue with a salient outcome (Lutas et al., 2019; Sengupta et al., 2018). Incorrect nose pokes that triggered a timeout did not elicit a different response in principal cell activity compared to before timeouts were incorporated (**Fig. S3.2 B-F**).

**Fig. 3.**
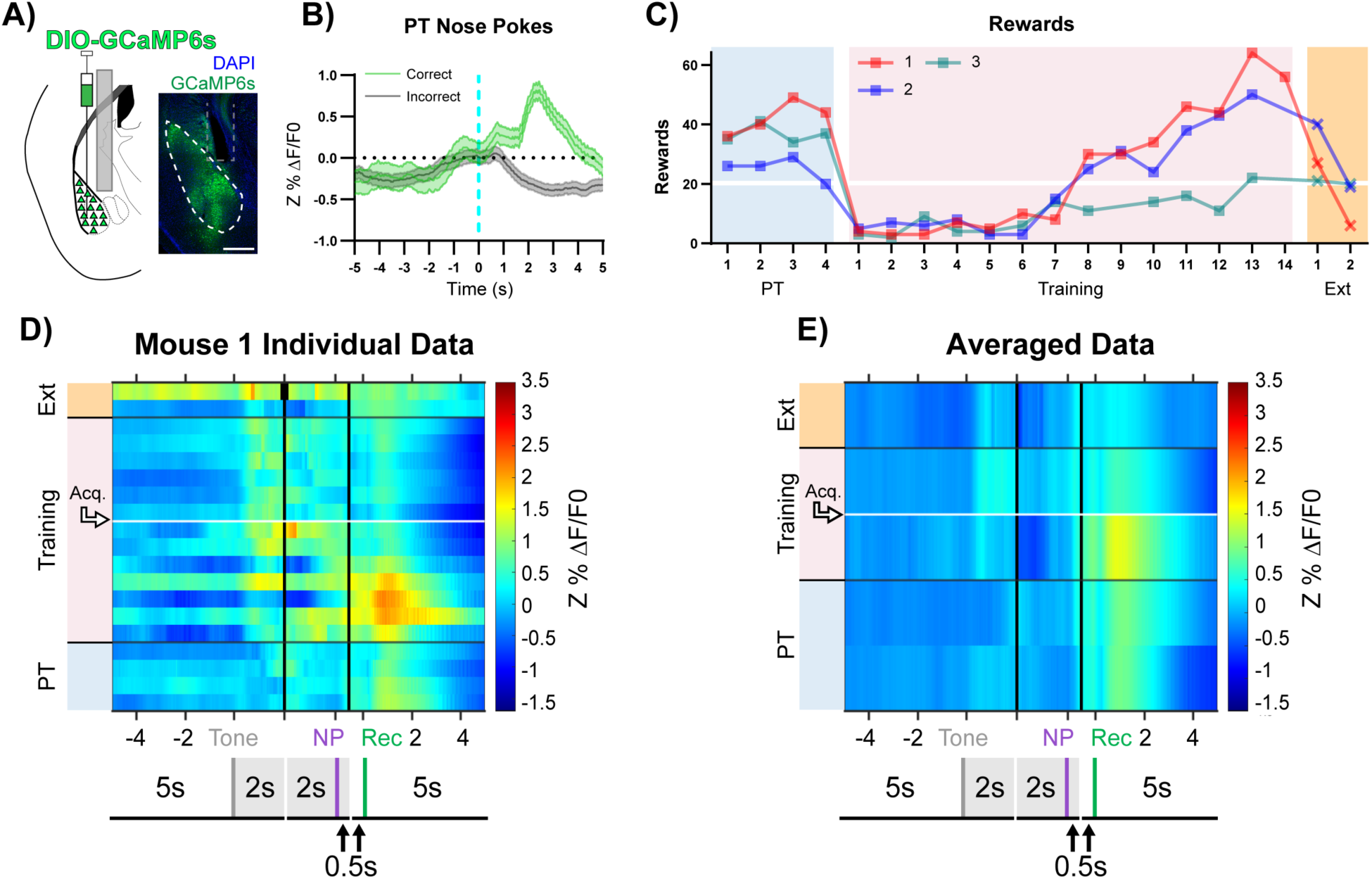
BLA Principal Neuron Activity Aligns to Reward Retrieval and Cue-Reward Learning A) <u>Diagram and example of injection and fiber placement sites in the BLA for recording from CaMKIIα-Cre mice expressing a fluorescent calcium indicator (DIO-GCaMP6s).</u> Left: Diagram of injection and fiber placement. Right: Representative coronal brain slice with fiber tip and GCaMP6s expression. White dashed line: BLA outline. Grey dashed rectangle: fiber tract. Blue: DAPI, Green: GCaMP6s. Scale 500 µm. Individual fiber placements are shown in **Fig. S3.1A**. B) <u>Fluorescence traces from BLA of GCaMP6s-expressing CaMKIIα-Cre mice</u>. During the last day PT, (data shown for Mouse 1) correct nose pokes (green line) were followed by a modest rise in BLA principal cell activity that increased steeply following receptacle entry (**Fig. S3.1B**) while incorrect nose pokes (grey line) were followed by a persistent decrease in activity. Mean ± SEM, correct (n = 44), incorrect (n = 141). Signal and reference channels (%ΔF/F0) during nose pokes are shown in **Fig. S3.1C-D**. Incorrect nose pokes on last day of PT vs Training Day 1 shown in **Fig. S3.2B.** C) <u>Behavioral responding of CaMKIIα-Cre mice expressing GCaMP6s in BLA</u>. Individual mice acquired the task at different rates as measured by rewards earned. Horizontal white line: acquisition threshold, when a mouse began to earn ∼20 rewards consistently in Training. Incorrect nose pokes shown in **Fig. S3.2A**. D) <u>Heatmap of BLA principal cell activity (Mouse 1) across all training phases, aligned to tone onset (Tone), correct nose poke (NP), and receptacle entry (Rec)</u>. Each row is the average of rewarded trials across a training session. White horizontal line: Day acquisition threshold met, as determined by rewards earned. Black horizontal lines: divisions between training phases. Black vertical lines: divisions between breaks in time to allow for variable latencies in tone onset, correct nose poke, and receptacle entry. Blanks in the heatmaps indicate time bins added for alignment. Individual data for mice 2-3 in **Fig. S3.1E-F**. Incorrect nose pokes heatmaps for individual mice shown in **Fig S3.2C-E.** E) <u>Heatmap of BLA principal cell activity averaged across mice</u>. Signal aligned as in D) with a selection of key days shown, from bottom to top: PT Day 1, PT Day 4, Training Day 3, Acquisition day (white horizontal line), Last Extinction Day. Black horizontal lines: divisions between training phases. Black vertical lines: divisions between breaks in time to allow for variable latencies in tone onset, correct nose poke, and receptacle entry. Incorrect nose poke heatmaps averaged across mice shown in **Fig. S3.2F.**

### Stimulation of cholinergic terminals in BLA improves cue-reward learning

Since ACh released by NBM-BLA terminals during Training shifted to tone onset during acquisition of cue-reward learning **(Fig. 2E, J**), we hypothesized that ACh may potentiate learning the cue-reward contingency. We therefore tested whether increasing ACh release in BLA during learning could alter cue-reward learning by injecting AAV carrying a Cre-recombinase-dependent channelrhodopsin-EYFP (AAV-DIO-ChR2-EYFP) construct bilaterally into the NBM of ChAT-IRES-Cre transgenic mice and placing fibers over the BLAs to optically stimulate cholinergic terminals originating from the NBM selectively (**Fig. 4A + Fig. S4.1**). Optical control over ChAT^+^ NBM cells was verified by *ex vivo* slice recordings (**Fig. 4B**). After shaping, ChAT^+^ NBM-BLA terminals were stimulated via bilateral optical fibers triggered by a correct nose poke throughout both Pre-Training (**Fig. 4C**) and Training (**Fig. 4D**). Stimulation occurred during at least a portion of all three components of a rewarded trial: tone, correct nose poke, and reward retrieval, since these events were often separated by short latencies.

**Fig. 4.**
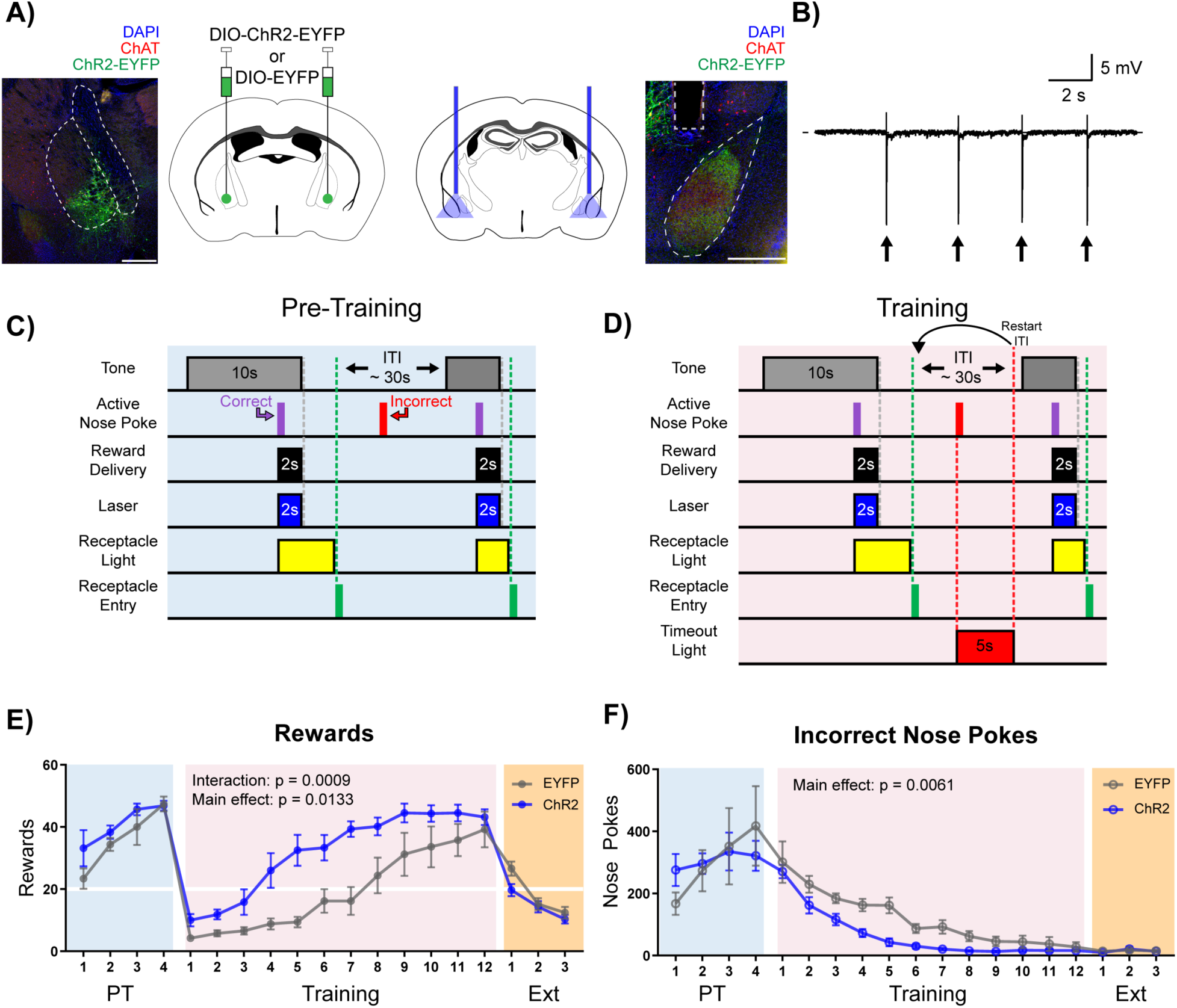
Stimulation of Cholinergic Terminal Fibers in the BLA Enhances Cue-Reward Learning. A) Schematic of optical stimulation of ChAT^+^ terminal fibers projecting to the BLA. Left: Bilateral AAV injection into the NBM of ChAT-IRES-Cre mice to gain optical control over ChAT^+^ NBM cells and representative coronal brain slice showing ChR2-EYFP expression. White dashed lines: internal capsule and globus pallidus outlines. Blue: DAPI, red: ChAT, green: ChR2-EYFP. Scale: 500 µm, individual injection sites shown in **Fig. S4.1A** and separate channels shown in **Fig. S4.1B**. Right: Bilateral optical fiber implantation above BLA to stimulate BLA-projecting ChAT^+^ NBM cells. Representative coronal brain slice showing ChR2-EFYP expression and fiber tip placement. Grey dashed rectangle: fiber tract. White dashed: BLA outline. Blue: DAPI, red: ChAT, green: ChR2-EYFP. Scale: 500 µm, individual fiber tip placements shown in **Fig. S4.1C** and separate channels shown in **Fig. S4.1D**. Injection sites and fiber tip placements for males from **Fig. S4.2C-F** shown in **S4.3A-B.** B) <u>Optical stimulation validation *via* local field potential recordings</u>. Extracellular recording of action potentials induced by optical stimulation of ChAT^+^ NBM cells expressing ChR2. Arrows indicate 60 ms laser pulse. <u>C-D) Details of the Cue-Reward Learning Paradigm</u> C) During Pre-Training, auditory tones were presented on a variable interval 30 schedule (VI30), during which an active nose poke (correct) yielded Ensure reward delivery and 2 sec of optical stimulation but there was no consequence for incorrect nose pokes (active nose pokes not during tone). D) Training was identical to Pre-Training, except incorrect nose pokes resulted in a 5 sec timeout, signaled by house light illumination, followed by a restarting of the ITI. E) <u>Behavioral performance in a cue-reward learning task improves with optical stimulation of</u> ChAT^+^ fibers in BLA. EYFP- and ChR2-expressing mice earn similar numbers of rewards during PT (blue shaded region). ChR2-expressing mice more rapidly earn significantly more rewards than EYFP-expressing mice during Training (pink shaded region). No significant differences were observed during extinction training (orange shaded region). Horizontal white line: acquisition threshold, when a mouse began to earn ∼20 rewards consistently in Training. Mean ± SEM, EYFP: n = 5, ChR2: n = 6. Individual data are shown in **Fig. S4.2A**. Data for males shown in **Fig. S4.2C,E.** F) <u>EYFP- and ChR2-expressing mice made similar numbers of incorrect nose pokes during Pre-Training</u>. ChR2-epxressing mice made significantly fewer incorrect nose pokes than EYFP-expressing mice in Training. No significant differences were observed during extinction training. Mean ± SEM, EYFP: n = 5, ChR2: n = 6. Individual data are shown in **Fig. S4.2B**. Data for males shown in **Fig. S4.2D,F.** Additional behavioral assays shown in **Fig. S4.4A-F**.

As seen in previous experiments, during the Pre-Training phase animals made a high number of nose poke responses over the course of each session, obtained most available rewards by the last day (**Fig. 4E + Fig. S4.2A**, blue shading), and committed a very high number of incorrect nose pokes (**Fig. 4F + Fig. S4.2B**, blue shading). There were no differences in rewards earned (main effect of group (EYFP vs. ChR2) in a two-way repeated-measures ANOVA, F (1, 9) = 1.733, p = 0.2205) or incorrect nose pokes (main effect of group (EYFP vs. ChR2) in a two-way repeated-measures ANOVA, F (1, 9) = 0.002433, p = 0.9617) between the EYFP control (n = 5) and ChR2 (n = 6) groups during the Pre-Training phase (**Fig. 4E-F + Fig. S4.2A-B**, blue shading), suggesting that increasing BLA ACh signaling was not sufficient to modify behavior during the Pre-Training phase of the task.

On Day 1 of the Training phase, all animals earned fewer rewards (**Fig. 4E + Fig. S4.2A**, pink shading) and incorrect nose pokes remained high (**Fig. 4F + Fig. S4.2B**, pink shading). As the animals learned that a nose poke occurring outside of the cued period resulted in a timeout, both control EYFP and ChR2 groups learned the contingency and improved their performance, resulting in acquisition of the cue-reward task (20 rewards earned). However, significant group differences emerged, such that ChR2 mice earned significantly more rewards than EYFP controls (**Fig. 4E + Fig. S4.2A**, pink shaded; main effect of group (EYFP vs. ChR2) in a two-way repeated-measures ANOVA, F (1, 9) = 9.434, p = 0.0133), and there was a significant Day x Group (EYFP vs. ChR2) interaction (two-way repeated-measures ANOVA, F (11, 99) = 3.210, p = 0.0009). ChR2 mice also made significantly fewer incorrect nose pokes than control mice (**Fig. 4F + Fig. S4.2B**, pink shaded; two-way repeated-measures ANOVA, F (1, 9) = 12.67, p = 0.0061), suggesting that the ChR2 group learned the tone-reward contingency more quickly than the EYFP group. EYFP mice were able to reach the same peak cue-reward performance as the ChR2 group only after 4-6 additional days of training. Once peak performance was achieved, there was no difference in extinction learning between the groups (main effect of group (EYFP vs. ChR2) in a two-way repeated-measures ANOVA, F (1, 9) = 2.293, p = 0.1643). While sex differences in the behavior were not formally tested side by side, an independent cohort of male mice (EYFP n = 7, ChR2 n = 7, **Fig. S4.3**) was tested to determine whether both male and female mice would respond to ACh stimulation, revealing similar trends during Training for rewards earned (**Fig. S4.2C,E**, pink shaded; two-way repeated-measures ANOVA, Group main effect (EYFP vs. ChR2): F (1, 12) = 3.636, p = 0.0808, Day x Group interaction: F (11, 132) = 3.033, p = 0.0012) and incorrect nose pokes (**Fig. S4.2D,F**, red shaded; two-way repeated-measures ANOVA, Group main effect (EYFP vs. ChR2): F (1, 12) = 4.925, p = 0.0465).

In order to determine if optical stimulation of NBM-BLA cholinergic terminals improved performance in the task by increasing the rewarding value of the outcome, rather than enhancing cue-reward learning by some other means, we allowed mice to nose poke for optical stimulation rather than for Ensure (**Fig. S4.4A**). There were no differences between the EYFP control and ChR2 groups (two-way repeated-measures ANOVA, F (1, 9) = 0.6653, p = 0.4357). We also tested whether NBM-BLA cholinergic terminal activation was reinforcing on its own by stimulating these terminals in a real-time place preference test. Mice were allowed to explore two similar compartments to determine baseline preference, and NBM-BLA cholinergic terminals were then stimulated in one of the two chambers to determine whether it increased time spent in the simulation-paired chamber. There was no difference between groups (**Fig. S4.4B**, main effect of group (EYFP vs. ChR2) in a two-way repeated-measures ANOVA, F (1, 9) = 0.1311, p = 0.7257) in place preference, confirming that optical activation of NBM-BLA cholinergic terminals is not innately rewarding. Stimulation of NBM-BLA cholinergic terminals also did not lead to changes in nose poke behavior in an uncued progressive ratio task (**Fig. S4.4C**, main effect of group (EYFP vs. ChR2) in a two-way repeated-measures ANOVA, F (1, 12) = 0.0009814, p = 0.975). Locomotor behavior was also not significantly affected by NBM-BLA cholinergic terminal activation (**Fig. S4.4D**, two-way repeated-measures ANOVA, F (1, 9) = 0.05804, p = 0.8150.) Finally, to determine whether there was any effect of NBM-BLA cholinergic terminal stimulation on preference for, or avoidance of, a stressful environment, mice were tested for changes in time spent in the dark or light side due to laser stimulation in the Light/Dark Box test, and there were no differences between the groups (**Fig. S4.4E-F**, unpaired t-tests, number of crosses: p = 0.3223; time in light side: p = 0.1565).

### Muscarinic, but not nicotinic, receptors are required for acquisition of the cue-reward **contingency**

ACh signals through multiple receptor subtypes, with rapid, ionotropic signaling mediated through stimulation of nAChRs, and metabotropic signaling mediated through stimulation of mAChRs (Picciotto et al., 2012). To determine which ACh receptors were involved in this cue-reward learning task, mice were injected intraperitoneally with saline (n = 8), mecamylamine (non-competitive nicotinic antagonist, Mec, n = 9), scopolamine (competitive muscarinic antagonist, Scop, n = 8), or a combination of both antagonists (Mec+Scop, n = 9) 30 min prior to Pre-Training and Training, during the same epochs of the task in which optical stimulation was administered (**Fig. 5A**). Like optical stimulation, blockade of ACh receptors during the Pre-Training phase of the task had no effect on rewards earned (**Fig. 5B + Fig. S5.1A**, blue shading, main effect of Group (antagonist) in a two-way repeated-measures ANOVA, F (3, 30) = 1.285, P=0.2973) or on the large number of incorrect nose pokes (**Fig. 5C + Fig. S5.1B**, blue shading, main effect of Group (antagonist) in a two-way repeated-measures ANOVA, F (3, 30) = 1.496, p = 0.2356). In contrast, blockade of muscarinic signaling abolished the ability of mice to learn the correct cue-reward contingency during the Training period (**Fig. 5B + Fig. S5.1A**, pink shading, two-way repeated-measures ANOVA, Antagonist main effect: F (3, 30) = 23.13, p < 0.0001, Day x Antagonist interaction: F (33, 330) = 10.79, p < 0.0001), with these mice maintaining high levels of incorrect nose pokes for the duration of Training compared to Saline and Mec treated mice (**Fig. 5C + Fig. S5.1B**, pink shading, main effect of Group (antagonist) in a two-way repeated-measures ANOVA, F (3, 30) = 25.64, p < 0.0001). Saline and Mec groups were not significantly different in any phase of the task, including across Extinction (**Fig. 5B-C + Fig. S5.1A-B**, orange shading, main effect of Group (antagonist) in a two-way repeated-measures ANOVA, F (1, 15) = 1.201, p = 0.2903). Consistent with the inability to acquire the cue-reward contingency, mice treated with Scop or Mec+Scop also obtained very few rewards during Extinction (**Fig. 5B + Fig. S5.1A**, orange shading). The antagonists had no effect on locomotion as measured by beam breaks (**Fig. S5.1C**) one-way ANOVA, F (3, 30) = 0.5074, p = 0.6802).

**Fig. 5.**
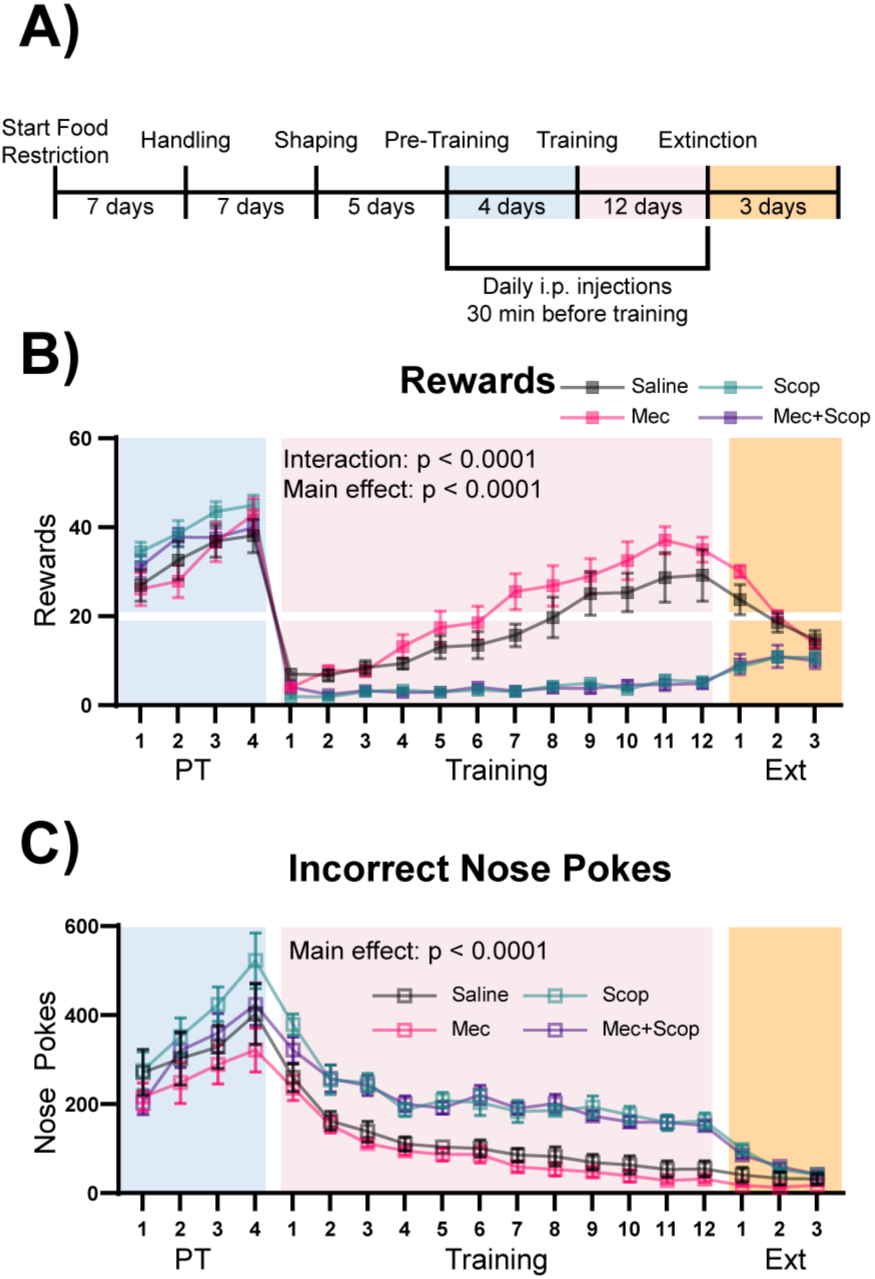
Muscarinic, but not Nicotinic, ACh Receptor Antagonism Prevents Learning of a Cue-Reward Contingency A) <u>Timeline of drug administration</u>. Saline or ACh receptor (AChR) antagonists were delivered, i.p., 30 min before PT and Training sessions, the same phases of the task as optical stimulation in Fig. 4. B) <u>Behavioral performance of mice administered AChR antagonists.</u> AChR antagonists had no significant effect on rewards earned during Pre-Training. Muscarinic AChR antagonism (Scop and Mec+Scop) resulted in significantly fewer rewards earned during Training. There was no significant difference between saline controls and those receiving the nicotinic AChR antagonist (Mec) during Training and mice extinguished responding at similar rates. Mean ± SEM Saline (n = 8), Mec (n = 9), Scop (n = 8), Mec+Scop (n = 9). Horizontal white line: acquisition threshold, when a mouse began to earn ∼20 rewards consistently in Training. Individual data are shown in Fig. S5.1A. C) <u>Incorrect nose pokes</u>. Incorrect nose poking was not affected by AChR antagonism during PT but Scop- and Scop+Mec-treated mice maintained high levels of incorrect nose pokes compared to Saline- and Mec-treated mice throughout Training. Mean ± SEM, Saline (n = 8), Mec (n = 9), Scop (n = 8), or Mec+Scop (n = 9). Individual data are shown in **Fig. S5.1B**. AChR antagonist locomotor test shown in **Fig. S5.1C**

### ACh-mediated accelerated cue-reward learning does not require contingent stimulation of ChAT^+^ NBM terminals in the BLA

Acetylcholine is often thought of as a neuromodulator (Picciotto et al., 2012), and the window for cholinergic effects on synaptic plasticity varies across ACh receptor subtypes (Gu & Yakel, 2011). It is therefore possible that ACh signaling may result in intracellular signaling changes that outlast the cue presentation window. In order to determine if the effect of NBM-BLA stimulation is dependent upon the timing of correct nose poke and laser stimulation contingency, we repeated the experiment in an independent cohort of mice with an additional yoked, non-contingent ChR2 group that received the same number of stimulation trains as the contingent ChR2 group, but in which light stimulation was explicitly unpaired with task events (**Fig. 6A + Fig. S6.1**). As in the previous experiment, there were no differences between the EYFP control (n = 6) and stimulation groups (contingent ChR2 n = 5 and Yoked non-contingent ChR2 n = 5) during Pre-Training (**Fig. 6B-C + Fig. S6.2 A-B**, blue shading; main effect of group (EYFP vs. contingent ChR2 vs. Yoked non-contingent ChR2) two-way repeated-measures ANOVAs; rewards earned: F (2, 13) = 0.7008, p = 0.5140; incorrect nose pokes: F (2, 13) = 0.3906, p = 0.6843). However, the Yoked non-contingent ChR2 group was not significantly different from the contingent ChR2 group during the Training period with respect to number of rewards earned (two-way repeated-measures ANOVA, F (1, 8) = 0.09147, p = 0.7700) or incorrect nose pokes (two-way repeated-measures ANOVA, F (1, 8) = 0.3681, p = 0.5609), but both ChR2 groups were significantly better than the EYFP control group (**Fig. 6B-C + Fig. S6.2 A-B**, pink shading; two-way repeated-measures ANOVAs; rewards earned: Group (EYFP vs. contingent ChR2 vs. Yoked ChR2) main effect: F (2, 13) = 7.254, p = 0.0077; Day x Group interaction: F (22, 143) = 1.861, p = 0.0164. Incorrect nose pokes: Group main effect: F (2, 13) = 4.884, p = 0.0262.). These results demonstrate that ACh release does not have to be time-locked to the cue, nose poke, or reward retrieval to improve performance of the task, suggesting that ACh may alter the threshold for neuronal plasticity for cue-reward pairing over a much longer timescale than might be expected based on results from the ACh3.0 recording and NBM-BLA recordings, which could be consistent with the involvement of mAChR signaling in this effect. As in the previous experiment, once all groups reached criterion for acquisition of the cue-reward contingency, there were no differences between any of the groups during Extinction (**Fig. 6B-C + Fig. S6.2 A-B**, orange shaded; two-way repeated-measures ANOVA, F (2, 13) = 0.04229, p = 0.9587).

**Fig. 6.**
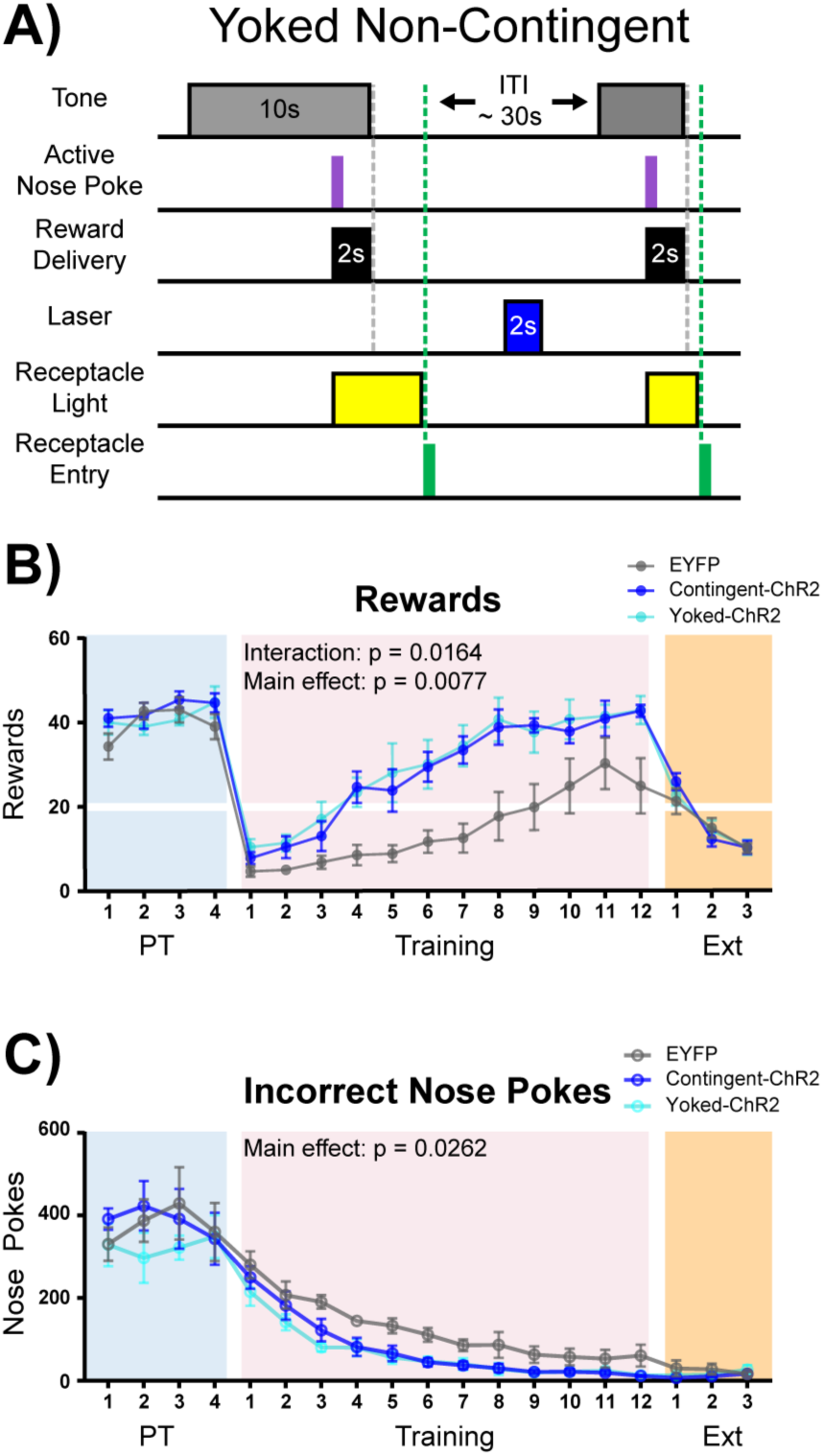
Non-Contingent Stimulation of ChAT^+^ NBM-BLA Terminals is Sufficient to Enhance Cue-Reward Learning A) <u>Experimental details of laser stimulation in non-contingent yoked mice.</u> Yoked non-contingent ChR2-expressing mice received the same number of light stimulations as contingent ChR2-expressing mice, but stimulation was only given during the ITI, when Yoked mice had not made a response within 2 sec. Injection sites and fiber placements are shown in **Fig. S6.1 A-B**. B) <u>Yoked non-contingent NBM-BLA optical stimulation also improves behavioral performance in cue-reward learning task.</u> There was no significant difference in the number of rewards earned between EYFP (n = 6), Contingent-ChR2 (n = 5), or Yoked-ChR2 (n = 5) mice during Pre-Training. Contingent- and Yoked-ChR2-expressing mice more rapidly earned significantly more rewards during Training than EYFP-expressing mice. No differences were observed between groups during extinction training. Mean ± SEM EYFP: n = 6, contingent-ChR2: n = 5, Yoked-ChR2: n = 5. Horizontal white line: acquisition threshold, when a mouse began to earn ∼20 rewards consistently in Training. Individual data are shown in **Fig. S6.2A**. C) <u>Incorrect nose pokes</u>. There was no significant difference in the number of incorrect nose pokes between groups during Pre-Training. Contingent- and Yoked-ChR2-expressing mice made significantly fewer incorrect nose pokes during Training than EYFP-expressing mice. No differences between groups were observed during extinction training. Mean ± SEM EYFP: n = 6, ChR2: n = 5, Yoked: n = 5. Individual data are shown in **Fig. S6.2B**.

## Discussion

It is increasingly recognized that the BLA is involved in learning to predict both positive and negative outcomes from previously neutral cues (Cador et al., 1989; Janak & Tye, 2015; LeDoux et al., 1990). Cholinergic cells in the basal forebrain complex fire in response to both positive and negative reinforcement (Hangya et al., 2015). The results shown here indicate that ACh signaling in the BLA is intimately involved in cue-reward learning. Endogenous ACh is released in the BLA in response to salient events in the task, and ACh dynamics evolved as the subject formed associations between stimuli and reward. While the pattern of ACh signaling in the BLA may seem reminiscent of how dopamine neurons encode reward prediction errors as measured in other brain areas (Schultz et al., 1997), the current results suggest that ACh release in the BLA may instead be involved in signaling a combination of salience and novelty. ACh release and NBM-BLA activity increased following correct nose poke and, around the time that animals acquired the cue-reward task, following tone onset. However, earlier in training, incorrect nose pokes that resulted in a timeout were also followed by ACh release, although this was lower in magnitude. Further, stimulating NBM-BLA cholinergic terminals during learning enhanced behavioral performance, but was not intrinsically rewarding on its own and did not support responding for the tone alone. Although ACh was released in the BLA at discrete points during the task, the effects of heightened BLA ACh signaling were relatively long lasting, since it was not necessary for stimulation to be time-locked to cue presentation or reward retrieval to enhance behavioral performance. Thus, cholinergic inputs from the basal forebrain complex to the BLA are a key component of the circuitry that links salient events to previously neutral stimuli in the environment and uses those neutral cues to predict future rewarded outcomes.

### BLA ACh signaling and principal cell activity are related to cue-reward learning

We have shown that ACh release in the BLA is coincident with the stimulus that was most salient to the animal at each phase of the task. Use of the fluorescent ACh sensor was essential in determining these dynamics. Previous microdialysis studies have shown that ACh is released in response to positive, negative, or surprising stimuli, but this technique is limited by relatively long timescales (minutes) and cannot be used to determine when cholinergic transients align to given events in an appetitive learning task and how they evolve over time (Sarter & Lustig, 2020). In this cue-reward learning paradigm, when there was no consequence for incorrect nose-poking (Pre-Training phase), animals learned to perform a very high number of nose pokes and received a large number of rewards, and BLA ACh signaling peaked following correct nose pokes. Both the behavioral response (nose poking that was not contingent with the tone) and the ACh response (linked to the correct nose poke) suggest that the animals were not attending to the tone during the Pre-Training phase of the task, but rather were attending to the cues associated with reward delivery, such as the reward light or the sound of the pump that delivered the reward. Consistent with this possibility, in the next phase of the task when mice received a timeout for responding if the tone was not presented, performance of all groups dropped dramatically. Interestingly, in the early Training sessions, ACh release shifted to reward retrieval, likely because this was the most salient aspect of the task when the majority of nose pokes performed did not result in reward. Finally, as mice acquired the contingency between tone and reward availability, the tone also began to elicit ACh release in the BLA, suggesting that mice learned that the tone is a salient event predicting reward availability. Since there are multiple sources of ACh input to the BLA, it was important to determine whether NBM cholinergic neurons were active during the periods when ACh levels were high (Woolf, 1991). Recordings from cholinergic NBM-BLA terminal fibers showed similar dynamics to ACh measurements, suggesting that the NBM is a primary source of ACh across the phases of cue-reward learning.

Perhaps the most well-known example of dynamic responding related to learning cue-reward contingencies and encoding of reward prediction errors is the firing of dopaminergic neurons of the ventral tegmental area (VTA; Schultz, 1998). After sufficient pairings, dopaminergic neurons will fire in response to the cue that predicts the reward, and no longer to the rewarding outcome, which corresponds with behavioral changes that indicate an association has been formed between conditioned stimuli (CS) and unconditioned stimuli (US). Plasticity related to learning has also been observed in cholinergic neurons in the basal forebrain complex during aversive trace conditioning, such that after several training days, neuronal activity spans the delay between CS and US (Guo et al., 2019). Additionally, a recent study suggested that ACh may signal a valence-free reinforcement prediction error (Sturgill et al., 2020). Future studies on the selective inputs to NBM to BLA cholinergic neurons would be of interest to identify the links between brain areas involved in prediction error coding.

We found that BLA principal cells were most reliably activated following reward retrieval before contingency acquisition (both when they were receiving several rewards but no timeouts in Pre-Training and few rewards early in Training). Similar to the recording of ACh levels, after acquisition, the tone began to elicit an increase in BLA principal cell population activity. However, activity of principal neurons differed from ACh signaling in the BLA in important ways. ACh was released in response to the salient events in the task that were best able to predict reward delivery or availability. In contrast, the activity of BLA principal neurons was not tightly time-locked to correct nose poking, and instead followed reward retrieval until acquisition, when activity increased in response to tone onset. The divergent dynamics of ACh release and principal neuron activity underscores that ACh’s role in the BLA is to modulate, rather than drive, the activity of principal neurons, and therefore may alter dynamics of the network through selective engagement of different populations of GABA interneurons (Unal et al., 2015).

### Increasing BLA acetylcholine levels enhances cue-reward learning

Neuronal activity and plasticity in the BLA is required for both acquisition of appetitive learning (conditioned reinforcement) and fear conditioning, however the inputs that increase activity in the structure during salient events likely come from many brain areas (McKernan & Shinnick-Gallagher, 1997; Rogan et al., 1997; Tye et al., 2008). In particular, dopaminergic inputs to the BLA are important for acquisition of conditioned reinforcement and for linking the rewarding properties of addictive drugs to cues that predict their availability (Cador et al., 1989). Our results indicate that ACh is a critical neuromodulator upstream of the BLA that is responsive to salient events, such as reward availability, motor actions that elicit reward, and cues that predict reward. We show here that increasing endogenous ACh signaling in the BLA caused mice to perform significantly better than controls in an appetitive cued-learning task. Heightened ACh release during learning of a cue-action-reward contingency led to fewer incorrect responses and increased acquisition rate in both female and male mice. The optical stimulation was triggered by correct nose poke, thus the cholinergic NBM-BLA terminal fiber stimulation overlapped with all three salient events: tone, nose poke, and reward retrieval, since the tone terminated 2 sec after correct nose poke. Therefore, the initial optical stimulation of ACh release coincided with the tone and correct nose poke from the beginning of training in ChR2 mice, approximating the ACh signature in mice that had already acquired the cue-reward contingency. We hypothesize that it was this premature increase in ACh levels at the time of cue presentation that was important in allowing the animals to learn the contingency earlier.

It is possible that ACh increased learning by increasing the intensity of the reward, potentiating the learned association, improving discrimination, or a combination of these phenomena. However, increasing ACh release in the BLA was not inherently rewarding, because it did not support self-stimulation or real-time place preference. This is at odds with a recent study that found stimulation of NBM-BLA cholinergic terminals could induce a type of place-preference and modest self-stimulation (Aitta-aho et al., 2018). It is possible that slight differences in targeting of ChR2 infusion or differences in the behavioral paradigm could be responsible for the lack of direct rewarding effects of optical ChAT terminal stimulation in the current study. Other recent work (Jiang et al., 2016) has demonstrated that stimulating this NBM-BLA cholinergic pathway is sufficient to strengthen cued aversive memory, suggesting that the effect of ACh in the BLA may not be inherently rewarding or punishing, but instead potentiates plasticity in the BLA, allowing learning of cue-outcome contingencies. Similarly, it is possible that ACh alters motor activity. However, there were no effects of optical stimulation on locomotion or responding in the inactive nose poke port. In addition, during the Pre-Training phase when there was no consequence for incorrect nose pokes, all groups earned the same number of rewards, regardless of optical stimulation or pharmacological blockade of ACh receptors, suggesting that ACh is not involved in the motor aspects of the task or the value of the reward. Indeed, differences emerged only during the Training phase, when attention to the tone was critical to earn rewards. Further, incorrect nose poking remained high for mice administered scopolamine. This suggests that scopolamine-treated animals were seeking the reward, as in the shaping and Pre-Training phases of training, but were unable to learn that they should only nose poke in response to the tone.

Cell-type-specific expression of AChRs and activity-dependent effects place cholinergic signaling at a prime position to shape BLA activity during learning. For instance, late-firing interneurons in the BLA exhibit nAChR-dependent EPSP’s when no effect is seen on fast-spiking interneurons, while principal neurons can be either excited or inhibited through mAChRs, depending on activity level of the neuron at the time of cholinergic stimulation (Unal et al., 2015). BLA mAChRs can support persistent firing in principal neurons and can be important for the expression of conditioned place preference behavior, as well as trace fear conditioning (Baysinger et al., 2012; Egorov et al., 2006; McIntyre et al., 1998). Similar to studies of trace fear conditioning, in which activity of the network over a delay period must be maintained, we found that metabotropic (mAChRs) but not ionotropic (nAChRs) ACh receptors were required for learning the contingency of this cue-reward task. The timing of cholinergic signaling is a critical factor in the induction of synaptic plasticity in other brain regions, so we hypothesized that the enhancement of cue-reward learning observed might be dependent upon when NBM-BLA terminal fibers were stimulated with respect to tone presentation and/or behavioral responses (Gu & Yakel, 2011). However, we found that heightened ACh signaling in the BLA improved behavioral performance even when stimulations were explicitly unpaired with the cue or correct nose poking. This suggests that the effect of increased cholinergic signaling in the BLA is long lasting, and that stimulation across a learning session is sufficient to potentiate synaptic events linking the cue to a salient outcome. Coupled with pharmacological evidence demonstrating that muscarinic signaling is necessary for reward learning in this task, this time course suggests the involvement of metabotropic signaling downstream of muscarinic receptors that outlasts the initial cholinergic stimulation.

To conclude, the abundant ACh input to the BLA results in ACh release in response to stimuli that predict reward in a learned cue-reward task. Mimicking this increase in cholinergic signaling results in accelerated learning of the cue-reward contingency. These findings are consistent with the hypothesis that ACh is a neuromodulator that is released in response to salient stimuli and suggests that ACh signaling may enhance neuronal plasticity in the BLA network, leading to accelerated cue-reward learning.

## Materials and Methods

### Animals

All procedures were approved by the Yale University Institutional Animal Care & Use Committee in compliance with the National Institute of Health’s Guide for the Care and Use of Laboratory Animals. Experiments were performed in mice of both sexes, in keeping with the NIH policy of including sex as a biological variable. Sex of mice in behavioral graphs is indicated by circles for females and squares for males.

Female and male heterozygous mice with Cre recombinase knocked into the choline acetyltransferase (ChAT) gene (ChAT-IRES-Cre, B6;129S6-Chattm2(cre)Lowl/J, Stock number: 006410; Jackson Laboratory, Bar Harbor, ME) were bred in house by mating ChAT-IRES-Cre, B6;129S6-Chattm2(cre)Lowl/J with C57BL6/J mice. CaMKIIα-Cre mice obtained from Ronald Duman (Casanova et al., 2001; Wohleb et al., 2016) were bred in house as above. C57BL6/J mice were obtained from The Jackson Laboratory at 6-10 weeks of age, and tested at 5-7 months of age, following at least one week of acclimation. All mice were maintained in a temperature-controlled animal facility on a 12-hour light/dark cycle (lights on at 7:00 AM). Mice were group housed 3-5 per cage and provided with *ad libitum* food and water until undergoing behavioral testing. Mice were single housed 1-3 weeks before surgery to facilitate food restriction and body weight maintenance.

### Surgical procedures

Surgical procedures for behavior were performed in fully adult mice at 4-6 months of age, age-matched across conditions. For viral infusion and fiber implantation, mice were anesthetized using isoflurane (induced at 4%, maintained at 1.5-2%) and secured in a stereotactic apparatus (David Kopf Instruments, Tujunga, CA). The skull was exposed using a scalpel and Bregma was determined using the syringe needle tip (2 µL Hamilton Neuros syringe, 30 gauge needle, flat tip; Reno, NV).

For fiber photometry surgeries, either 0.4 µL of AAV9 hSyn-ACh3.0 (Vigene Biosciences Inc.) to measure BLA ACh levels (**Fig. 2A-E + S2.1-S2.2**) or 0.5 µL of AAV1 Syn-FLEX-GCaMP6s-WPRE-SV40 (Addgene, Watertown, MA) to measure BLA principal cell calcium dynamics (**Fig. 3 + S3.1-S3.2**) was delivered unilaterally to the BLA (A/P; −1.34 mm, M/L + 2.65 mm, D/V −4.6 mm, relative to Bregma) of ChAT-IRES-Cre or CaMKIIα-Cre mice, respectively, at a rate of 0.1 µL/min. The needle was allowed to remain at the infusion site for 5 min before and 5 min after injection. A mono fiber-optic cannula (1.25 mm outer diameter metal ferrule; 6 mm long, 400 µm core diameter/430 µm outer diameter, 0.48 numerical aperture (NA), hard polymer cladding outer layer cannula; Doric Lenses, Quebec City, Quebec, Canada) was implanted above the BLA (A/P; −1.34 mm, M/L + 2.65 mm, D/V −4.25 mm) and affixed to the skull using opaque dental cement (Parkell Inc., Edgewood, NY). Cholinergic NBM-BLA terminal fiber calcium dynamic recording (**Fig. 2F-J + S2.3-S2.4**) surgeries were performed as above except AAV1-Syn-FLEX-jGCaMP7s-WPRE (Addgene) was infused unilaterally into the NBM (A/P: - 0.7 mm, M/L - 1.75 mm, D/V – 4.5 mm) of ChAT-IRES-Cre mice, with the optical fiber being placed above the ipsilateral BLA. The jRCaMP1b + ACh3.0 surgeries to simultaneously measure cholinergic NBM-BLA terminal fiber calcium dynamics and BLA ACh levels (**Fig. S2.5**) consisted of both the NBM and BLA infusions above, except AAV1 Syn-FLEX-NES-jRCaMP1b-WPRE-SV40 (Addgene) was infused the NBM of ChAT-IRES-Cre mice. The RCaMP sham mouse (**Fig. S2.5E,H**) was a wild-type littermate and thus had no jRCaMP1b expression.

Mice were allowed to recover in a cage without bedding with a microwavable heating pad underneath it until recovery before being returned to home cage. For two days following surgery, mice received 5 mg/Kg Rimadyl i.p (Zoetis Inc., Kalamazoo, MI) as postoperative care.

For optical stimulation experiments (**Fig. 4,6 + Fig. S4.1-S4.4 + S6.1-S6.2**), surgeries were performed as above except as follows: 0.5 µL of control vector (AAV2 Ef1a-DIO-EYFP) or channelrhodopsin (AAV2 Ef1a-DIO-hChR2(H134R)-EYFP; University of North Carolina Gene Therapy Center Vector Core, Chapel Hill, NC) was delivered bilaterally into the NBM (A/P: - 0.7 mm, M/L ± 1.75 mm, D/V – 4.5 mm) of ChAT-IRES-Cre mice. Mono fiber-optic cannulas (1.25 mm outer diameter zirconia ferrule; 5 mm long, 200 µm core diameter/245 µm outer diameter, 0.37 NA, polyimide buffer outer layer cannula; Doric Lenses) were inserted bilaterally above the basolateral amygdala (BLA, A/P; −1.22 mm, M/L ± 2.75 mm, D/V −4.25 mm). Mice were randomly assigned to EYFP or ChR2 groups, controlling for average group age.

For *ex vivo* electrophysiology experiments (**Fig. 4B**), the NBM was injected with DIO-ChR2-EYFP as described above, except mice were 8 weeks of age. The coronal brain slices containing the NBM were prepared after 2-4 weeks of expression. Briefly, mice were anesthetized with 1X Fatal-Plus (Vortech Pharmaceuticals, Dearborn, MI) and were perfused through their circulatory systems to cool down the brain with an ice-cold (4°C) and oxygenated cutting solution containing (mM): sucrose 220, KCl 2.5, NaH2PO4 1.23, NaHCO3 26, CaCl21, MgCl2 6 and glucose 10 (pH 7.3 with NaOH). Mice were then decapitated with a guillotine immediately; the brain was removed and immersed in the ice-cold (4°C) and oxygenated cutting solution to trim to a small tissue block containing the NBM. Coronal slices (300 µm thick) were prepared with a Leica vibratome (Leica Biosystems Inc., Buffalo Grove, IL) after the tissue block was glued on the vibratome stage with Loctite 404 instant adhesive (Henkel Adhesive Technologies, Düsseldorf, Germany). After preparation, slices were maintained at room temperature (23-25 C°) in the storage chamber in the artificial cerebrospinal fluid (ACSF) (bubbled with 5% CO2 and 95% O2) containing (in mM): NaCl 124, KCl 3, CaCl2 2, MgCl2 2, NaH2PO4 1.23, NaHCO3 26, glucose 10 (pH 7.4 with NaOH) for recovery and storage. Slices were transferred to the recording chamber and constantly perfused with ACSF with a perfusion rate of 2 ml/min at a temperature of 33 oC for electrophysiological experiments. Cell-attached extracellular recording of action potentials was performed by attaching a glass micropipette filled with ACSF on EYFP-expressing cholinergic neurons with an input resistance of 10-20 MΩ under current clamp. Blue light (488 nm) pulse (60 ms) was applied to the recorded cells through an Olympus BX51WI microscope (Olympus, Waltham, MA) under the control of the Sutter filter wheel shutter controller (Lambda 10-2, Sutter Instrument, Novato, CA). All data were sampled at 3-10 kHz, filtered at 3 kHz and analyzed with an Apple Macintosh computer using Axograph X (AxoGraph). Events of field action potentials were detected and analyzed with an algorithm in Axograph X as reported previously (Rao et al., 2008).

### Behavioral Testing

#### Habituation

One week after surgery, mice were weighed daily and given sufficient food (2018S standard chow, Envigo, Madison, WI) to maintain 85% free-feeding body weight. All behavioral tests were performed during the light cycle. Mice were allowed to acclimate to the behavioral room for 30 min before testing and were returned to the animal colony after behavioral sessions ended.

Two weeks after surgery, mice were handled 3 min per day for 7 days in the behavioral room. Mice were given free access to the reward (Ensure^→^Plus Vanilla Nutrition Shake solution mixed with equal parts water (Ensure); Abbott Laboratories, Abbott Park, IL) in a 50 mL conical tube cap in their home cages on the last 3 days of handling to familiarize them to the novel solution. Mice were also habituated to patch cord attachment during the last 3 days of handling for optical stimulation and fiber photometry experiments. Immediately before training each day, a patch cord was connected to their optical fiber(s) via zirconia sleeve (s) (1.25 mm, Doric Lenses) before being placed in the behavioral chamber.

#### Operant Training

All operant training was carried out using Med Associates modular test chambers and accessories (ENV-307A; Med Associates Inc., Georgia, VT). For optical stimulation experiments, test chambers were housed in sound attenuating chambers (ENV-022M). Two nose poke ports (ENV-313-M) were placed on the left wall of the chamber and the reward receptacle (ENV-303LPHD-RL3) was placed on the right wall. The receptacle cup spout was connected to a 5 mL syringe filled with Ensure loaded in a single speed syringe pump (PHM-100). Nose pokes and receptacle entries were detected by infrared beam breaks. The tone generator (ENV-230) and speaker (ENV-224BM) were placed outside the test chamber, but within the sound attenuating chamber, to the left. The house light (used for timeout, ENV-315M) was placed on top of the tone generator to avoid snagging patch cords. Each chamber had a fan (ENV-025F28) running throughout the session for ventilation and white noise. Behavior chambers were connected to a computer running MEDPC IV to collect event frequency and timestamps. For optical stimulation experiments, a hole drilled in the top of the sound attenuating chambers allowed the patch cord to pass through. BLA ACh3.0 (**Fig. 2A-E**) and principal cell GCaMP6s (**Fig. 3**) fiber photometry recordings occurred in a darkened behavioral room outside of sound attenuating chambers due to steric constraints with rigid fiber photometry patch cords. Later behavioral chamber customization allowed NBM-BLA terminal fiber (**Fig. 2F-J**) and jRCaMP1b/ACh3.0 (**Fig. S2.5**) mice to be tested inside sound attenuating chambers. For fiber photometry experiments, a custom receptacle was 3D printed that extended the cup beyond the chamber wall to allow mice to retrieve the reward with more rigid patch cords. In addition, the modular test chamber lid was removed and the wall height was extended with 3D printed and laser cut acrylic panels to prevent escape. Each mouse was pseudo-randomly assigned to behavioral chamber when multiple chambers were used, counterbalancing for groups across boxes.

Three weeks after surgery, initial behavioral shaping consisted of one 35 min session of Free Reward to demonstrate the location of reward delivery; all other sessions were 30 mins. During Free Reward shaping, only the reward receptacle was accessible. After 5 min of habituation, Ensure (24 µL over 2 seconds) was delivered in the receptacle cup and a light was turned on above the receptacle. The receptacle light was turned off upon receptacle entry. The next phase of shaping, mice learned to nose poke to receive reward on a fixed-ratio one (FR1) schedule of reinforcement. Mice in experiments involving manipulations (optical stimulation and antagonist studies) were pseudo-randomly assigned to left or right active (reinforced) nose poke port. Mice in fiber photometry experiments were all assigned to right active port to minimize potential across subject variability. The inactive (unreinforced) port served as a locomotor control. During FR1 Shaping, each nose poke response into the active port resulted in receptacle light and reward delivery. After the mice reached criterion on FR1 Shaping (group average of 30 rewards for 2 consecutive days, usually 4-5 days), mice were advanced to the Pre-Training phase. This phase incorporated an auditory tone (2.5-5 kHz, ∼60 dB) that lasted for at most 10 seconds and signaled when active nose pokes would be rewarded. Only active nose pokes made during the 10 sec auditory tone (correct nose pokes) resulted in reward and receptacle light delivery. The tone co-terminated with Ensure delivery. During Pre-Training, there was no consequence for improper nose pokes, neither in the active port outside the tone (incorrect nose pokes) nor in the inactive port (inactive nose pokes). The number of inactive nose pokes were typically very low after shaping and were not included in analysis. After reward retrieval (receptacle entry following reward delivery) the receptacle light was turned off and the tone was presented again on a variable intertrial interval schedule with an average interval of 30 sec (VI 30), ranging from 10 to 50 sec (Ambroggi et al., 2008). After 4-5 days of tone training, mice progressed to the Training phase, which had the same contingency as Pre-Training except incorrect nose pokes resulted in a 5 sec timeout signaled by house light illumination, followed by a restarting of the previous intertrial interval. Extinction was identical to Training except no Ensure was delivered in response to correct nose pokes. In order to promote task acquisition, mice that were not increasing number of rewards earned reliably were moved to a VI 20 schedule after 9 days of VI 30 Training for BLA ACh3.0 or 6-7 days for BLA principal cell mice. The VI 20 schedule was only needed for the two groups that were trained outside of the sound attenuating chambers.

Between mice, excrement was removed from the chambers with a paper towel. At the end of the day chambers were cleaned with Rescue Disinfectant (Virox Animal Health, Oakville, Ontario, Canada) and Ensure syringe lines were flushed with water then air. Mice were excluded from analyses if a behavioral chamber malfunctioned (e.g. syringe pump failed) or they received the improper compound. Fiber photometry mice were excluded from analyses if they did not meet the acquisition criterion by the last day of Training.

#### Optical Stimulation

Optical stimulation was generated by a 473 nm diode-pumped solid-state continuous wave laser (Opto Engine LLC, Midvale, UT) controlled by a TTL adapter (SG-231, Med Associates Inc.). The laser was connected to a fiber optic rotary joint (Doric Lenses) via a mono fiber optic patch cord (200 µm core, 220 µm cladding, 0.53 NA, FC connectors; Doric Lenses). The rotary joint was suspended above the sound attenuating chamber with a connected branching fiber optic patch cord (200 µm core, 220 µm cladding, 0.53 NA, FC connector with metal ferrule; Doric Lenses) fed into the behavioral box. Laser power was adjusted to yield 10-12 mW of power at each fiber tip. The stimulation pattern was 25 ms pulses at 20 Hz for 2 seconds modified from parameters in (Jiang et al., 2016). Optical stimulation was only delivered during the Pre-Training and Training phases of the operant task. Both control (EYFP) and experimental (ChR2) groups received identical light delivery, and stimulation was triggered by a correct nose poke and co-terminated with the auditory tone and Ensure delivery. For the Yoked non-contingent experiment, the number of light stimulations was yoked to the concurrently running ChR2 mouse. The timing of the non-contingent yoked stimulation was explicitly unpaired with correct nose pokes or tones, and was held in queue until the mouse had not made a response in the last 2 sec, a tone was not going to be delivered within the next 2 sec, or at least 5 sec had passed since the mouse entered the receptacle after earning reward.

### Fiber Photometry

#### Acquisition

Fluorescent measurements of ACh and calcium levels were recorded using two Doric Lenses 1-site Fiber Photometry Systems: a standard 405/465 nm system and a 405/465/560 nm system. The standard 405/465 system was configured as follows: the fiber photometry console controlled the two connectorized LEDs (CLEDs, 405 nm modulated at 208.616 Hz and 465 nm modulated at 572.205 Hz) through the LED module driver. Each CLED was connected via attenuating patch cord to the five-port Fluorescence MiniCube (FMC5_AE(405)_AF(420-450)_E1(460-490)_F1(500-550)_S). A pigtailed fiber optic rotary joint was connected to the MiniCube and suspended above the behavioral chamber with a rotary joint holder in order to deliver and receive light through the implanted optical fiber. The other end of the rotary joint was connected to the mono fiber optic patch cord via M3 connector and attached with a zirconia sleeve to the implanted fiber optic as above. The F1 (500-550 nm) port of the MiniCube was connected to the photoreceiver (AC low mode, New Focus 2151 Visible Femtowatt Photoreceiver, New Focus, San Jose, CA) via a fiber optic adapter (Doric Lenses) that was finally connected back to the fiber photometry console through an analog port. The 405/465/560 nm system was set up similarly, except a 560 nm LED was incorporated (modulated at 333.786 Hz), a six-port MiniCube with two integrated photodetector heads was used (iFMC6_IE(400-410)_E1(460-490)_F1(500-540)_E2(555-570)_F2(580-680)_S), and Doric Fluorescence Detector Amplifiers were used (AC 1X or 10X mode, DFD_FOA_FC). A TTL adapter (SG-231, Med Associates Inc.) was connected to the digital input/output port to allow for timestamping when events occurred in the behavioral chamber. Signal was recorded using Doric Neuroscience Studio (V 5.3.3.14) via the Lock-In demodulation mode with a sampling rate of 12.0 kS/s. Data were decimated by a factor of 100 and saved as a comma-separated file.

#### Analysis

Preprocessing of raw data was performed using a modified version of a MATLAB (MathWorks, Natick, MA) script provided by Doric. The baseline fluorescence (F0) was calculated using a first order least mean squares regression over the ∼30 min recording session. Second order least mean squares regressions were used when photobleaching of the sensor was more pronounced, as in the case of NBM-BLA terminal fiber recordings. The change in fluorescence for a given timepoint (ΔF) was calculated as the difference between it and F0, divided by F0, which was multiplied by 100 to yield % ΔF/F0. The % ΔF/F0 was calculated independently for both the signal (465 nm) and reference (405 nm) channels to assess the degree of movement artifact. Since little movement artifact was observed in the recordings (**Fig. S2.1B-C, S2.3E-F, S3.1C-D**, tan lines), the signal % ΔF/F0 was analyzed alone. The % ΔF/F0 was z-scored to give the final Z % ΔF/F0 reported here. For the BLA principal cell recordings (**Fig. S3.1C-D)**, some mirroring of the signal channel observed in the reference channel. This is likely because 405 nm is not the “true” isosbestic point for GCaMP and we were instead measuring some changes in calcium-unbound GCaMP rather than calcium-insensitive GCaMP signal alone (Barnett et al., 2017; C. K. Kim et al., 2016; Sych et al., 2019). Graphs and heatmaps for averaged traces aligned to actions were based on licking bout epoch filtering code from TDT (Alachua, FL; link in code comments).

#### Heatmaps

Combined action heatmaps were generated in MATLAB (2019b) by analyzing data 5 sec preceding tone onset (rewarded trials only) to 5 sec after receptacle entry. Actions were aligned despite variable latencies by evenly splitting a maximum of 4 sec post-tone onset/pre-correct nose poke and 1 sec post-correct nose poke/pre-receptacle entry for each trial within a day. The resulting aligned trials were averaged to generate daily averages that made up the rows of the individual animal heatmaps. Blanks in the rows of heatmaps (black time bins) indicate time bins added for alignment, meaning that no trials for that day had a latency that stretched the entire window. Only rewarded trials where the mouse entered the receptacle within 5 sec after nose poke were analyzed. Full or partial training days were excluded from analysis if there were acquisition issues such as the patch cord losing contact with the fiber or behavioral apparatus malfunction. Lack of trials for analysis or recording issues led to missing rows of fiber photometry data in the heatmap despite having behavioral data, in which case these rows were skipped rather than adding entire blank rows. Due to individual differences in behavior, across-mouse average data was calculated by using a selection of days in which behavior was roughly similar or milestones such as first and last day of Pre-Training, first day earning 10 rewards in Training, first day crossing acquisition threshold (and maintaining afterward), last day of Training, last day of Extinction (with 4 or more rewarded trials that met analysis criteria). Additional days were included in across-mouse average heatmaps when possible. Incorrect nose poke heatmaps were generated by averaging signals for 5 sec before and 5 sec after incorrect nose pokes that were not preceded by an incorrect nose poke in the last 5 sec. The incorrect nose poke heatmaps averaged across mice were generated using the same selection of days as the combined action heatmaps for a given experiment.

### Pharmacology

Male wildtype C57BL/6J mice were injected i.p. 30 min prior to each Pre-Training and Training session with a volume of 10 mL/kg with the following compounds: 1X DPBS (Thermo Fisher Scientific, Waltham, MA), 1 mg/kg mecamylamine hydrochloride (Millipore Sigma, St. Louis, MO), 0.5 mg/kg (-) scopolamine hydrochloride (Millipore Sigma), or 1 mg/kg mecamylamine + 0.5 mg/kg scopolamine (**Fig. 5** + **Fig. S5.1**)

### Histology

After completion of behavioral experiments, animals were anesthetized with 1X Fatal-Plus (Vortech Pharmaceuticals). Once there was no response to toe-pinch, mice were transcardially perfused with 20 mL ice cold 1X DPBS followed by 20 mL 4% paraformaldehyde (PFA, Electron Microscopy Sciences, Hatfield, PA). Brains were extracted and post-fixed for at least 1 day in 4% PFA at 4°C and transferred to 30% sucrose (Millipore Sigma) for at least 1 day at 4°C. Brains were sliced 40 µm thick on a self-cooling microtome and stored in a 0.02% sodium azide (Millipore Sigma) PBS solution. Brain slices were washed in PBS, blocked for 2-3 hours (0.3% Triton X-100, American Bioanalytical, Canton, MA; 3% normal donkey serum, Jackson ImmunoResearch, West Grove, PA), then incubated overnight with primary antibodies (1:1000 + 1% normal donkey serum). Slices were then washed in PBS and incubated with secondary antibodies (1:1000) for 2 hours, washed, stained with DAPI for 5 min, washed, mounted, and coverslipped with Fluoromount-G (Electron Microscopy Sciences). All incubations were at room temperature. Microscope slides were imaged using a FLUOVIEW FV10i confocal microscope (Olympus). Injection sites and fiber placements were designated on modified Allen Mouse Brain Atlas figures (Lein et al., 2007). Mice were excluded from analyses if fluorescence was not observed at injection sites.

#### Antibodies used

Goat anti-ChAT (AB144P, Millipore Sigma)

Chicken anti-GFP (A-10262, Invitrogen, Carlsbad, CA)

Alexa Fluor 488 Donkey anti-Chicken (703-545-155, Jackson ImmunoResearch Inc.) Alexa Fluor 555 Donkey anti-Goat (A-21432, Invitrogen)

### Statistical Analyses

Operant behavioral data saved by MEDPC IV was transferred to Excel using MPC2XL. Data were organized in MATLAB and analyzed in Prism (V8.3.0, GraphPad Software, San Diego, CA). Differences between groups and interactions across days for Training were evaluated using Two-Way Repeated Measures ANOVAs. We computed the required sample size for a 90% power level with an alpha of 0.05 by estimating the control (EYFP) group mean would be 10 rewards and the mean experimental (ChR2) group would be 20 rewards with a standard deviation of 5. We utilized a power calculator for continuous outcomes of two independent samples, assuming a normal distribution. The result was 6 samples per group. Each manipulation experiment started with at least 6 mice were included in each group (*Sealed Envelope | Power calculator for continuous outcome superiority trial*, n.d.). In each experiment, each animal within a group served as a biological replicate. These studies did not include technical replicates. Masking was not applied during data acquisition but data analyses were semi-automated in MATLAB and performed blind to condition

## Supplemental Methods

### Cued Self-Stimulation

After Extinction, responding was reinstated in Training for 2 days. Then mice underwent a modified Training paradigm where correct nose pokes yielded only laser stimulation, without Ensure delivery.

### Real Time Place Preference

An empty, clear mouse cage (29.5 cm x 19 cm x 12.5 cm) had half of its floor covered in printer paper to provide a distinct floor texture. A video camera was placed above the cage and was connected to a computer running EthoVision XT (version 10.1.856, Noldus, Wageningen, Netherlands) to track the position of the mouse and deliver optical stimulation when the mouse was on the laser-paired side (via TTL pulse to OTPG_4 laser controller (Doric Lenses) connected to the laser; 20 Hz, 25 ms pulses). Mice were randomly assigned and counterbalanced to receive laser stimulation only on one side of the cage. Mice were allowed free access to either side for 15 min during a session. Baseline was established in the absence of optical stimulation on Day 1. Mice then received optical stimulation on Day 2 only when on the laser-paired side. Data are presented as percent time spent on the laser-paired side.

### Progressive Ratio testing

In the progressive ratio test, mice were given 60 min to nose poke for Ensure and 2 sec of optical stimulation on a progressive ratio schedule (escalations given below). Training Day escalation: 1, 2, 2, 2, 2, 3, 3, 3, 3, 3, 5, 5, 5, 5, 5, 8, 8, 8, 8, 8, 8, 11, 11, 11, 11, 11, 11, 15, 15, 15, 15, 15, 15, 22, 22, 22, 22, 22, 33, 33, 33, 33, 33, 44, 44, 44, 44, 44, 55, 66, 77, 88, 99, 133, 166, 199, 255, 313, 399, 499, 599, 777, 900,1222. Test Day escalation: 1, 2, 4, 6, 9, 12, 15, 20, 25, 32, 40, 50, 62, 77, 95, 118, 145, 178, 219, 268, 328, 402, 492, 603, 777, 900, 1222.

### Locomotor Activity

#### Optical Stimulation

Mice were placed in a square box (47 cm x 47 cm x 21 cm) for 20 min with a floor of filter paper that was changed between mice. During the 3^rd^ 5 min bin of the session, mice received optical stimulation (20 sec on/off, 20 Hz, 25 ms pulses). Locomotor activity was recorded via overhead camera and analyzed in 5 min bins with EthoVision.

#### Antagonists

Locomotor data was collected using an Accuscan Instruments (Columbus, Ohio) behavior monitoring system and software. Mice were individually tested in empty cages, with bedding and nesting material removed to prevent obstruction of infrared beams. Mice were injected (i.p.) with saline, mecamylamine (1 mg/kg, Sigma), scopolamine (0.5 mg/kg, Sigma), or mecamylamine+scopolamine (1 mg/kg and 0.5 mg/kg, respectively) 30 min before locomotor testing. Locomotion was monitored for 20 min using 13 photocells placed 4 cm apart to obtain an ambulatory activity count, consisting of the number of beam breaks recorded during a period of ambulatory activity (linear motion rather than quick, repetitive beam breaks associated with behaviors such as scratching and grooming).

### Light/Dark Box Exploration

A rectangular box was divided evenly into a light (clear top, illuminated by an 8W tube light) and dark (black walls, black top) side with a black walled divider in the middle with a small door. The lid and divider were modified to allow the optical fiber and patch cord to pass through freely. Mice were placed facing the corner on the light side furthest from the divider and the latency to crossing to the dark side was measured. The number of crosses and time spent on each side were measured for 6 min following the initial cross.

## Acknowledgements

These studies were supported by grants DA14241, DA037566, MH077681. LW, DT and PR were supported by NS022061, MH109104 from the National Institutes of Health, and by the intramural programs of NINDS and NIMH. X-BG was supported by DA046160. RBC was supported by T32-NS007224. We thank Samantha Sheppard for the use of her mouse illustration and animal care assistance and Nadia Jordan-Spasov for genotyping and laboratory help. Angela Lee, and Wenliang Zhou provided helpful input into experimental planning. Marcelo Dietrich, Eric Girardi, Usman Farooq, Onur Iyilikci, Sharif Kronemer, Matthew Pettus, and Zach Saltzman provided insightful discussion and assistance with analysis and figure design. Ralph DiLeone, Jane Taylor, and Hyojung Seo offered helpful discussion about experimental design and analysis. The support teams at Doric Lenses (Alex Côté and Olivier Dupont-Therrien) and Tucker-Davis Technologies provided discussion, analysis support, and MATLAB code assistance.

---

**Fig. S2.1.**
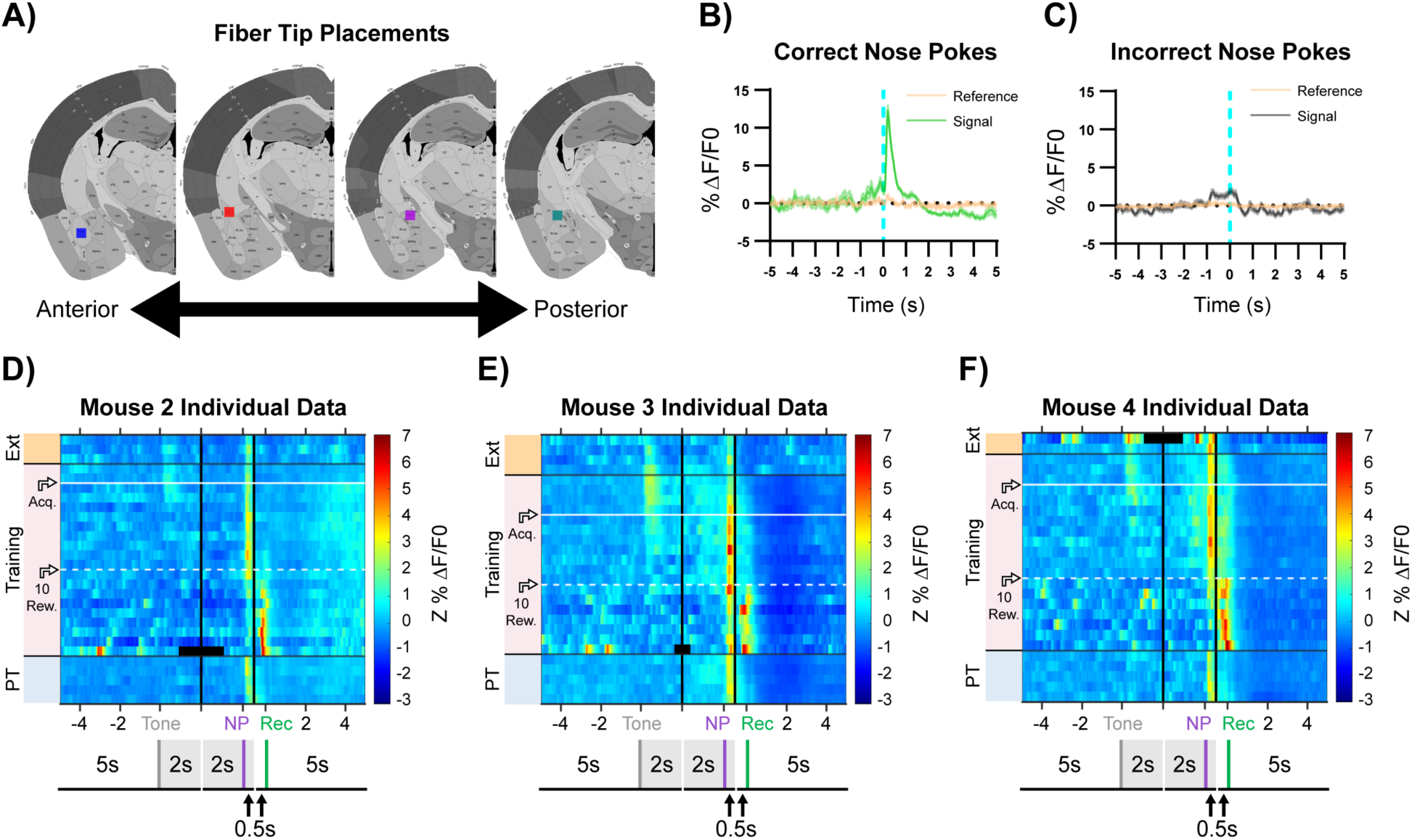
Supplemental Data for Fig. 2 A-E A) Squares indicate optical fiber tips for individual mice. 1 (red), 2 (blue), 3 (teal), 4 (purple). B) Increase in fluorescence (%ΔF/F0) following correct nose pokes is specific to the signal (465 nm, green) channel and is not observed in the reference channel (405 nm, tan). Data from Mouse 1 PT Day 5 as in Fig. 2C. Mean ± SEM, n = 24. C) Minimal increase in fluorescence (%ΔF/F0) following incorrect nose pokes. Signal (465 nm, grey) channel, reference channel (405 nm, tan). Data from Mouse 1 PT Day 5 as in Fig. 2C. Mean ± SEM, n = 58. D-F) Individual mouse data for mice 2-4 as shown in Fig. 2D. Dashed white horizontal line: first Training day earning 10 rewards (10 Rew.). White horizontal line: acquisition threshold (Acq.).

**Fig. S2.2.**
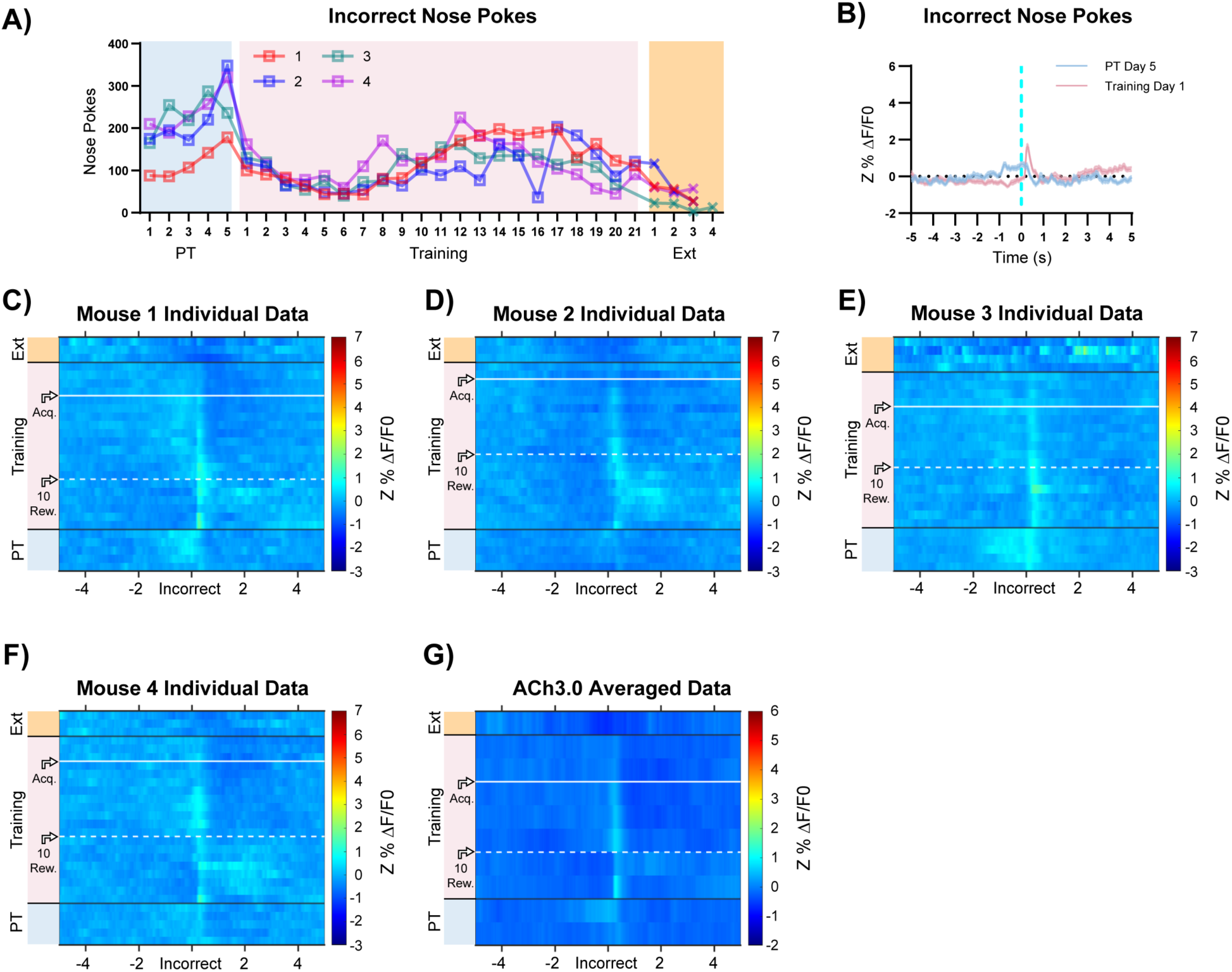
Supplemental Data for Fig. 2 A-E A) Incorrect nose poking of individual mice throughout training. B) Incorrect nose pokes that yield timeouts (Training Day 1, pink line, n = 66) result in increase in BLA ACh signaling but incorrect nose pokes before timeouts are introduced (PT Day 5, blue line, n = 58) do not. Data from Mouse 1 as in Fig. 2C, Mean ± SEM. C-F) Individual mouse heatmaps of BLA ACh signaling across all training phases, aligned to incorrect nose poke. Each row is the average of incorrect nose pokes that led to (or would have led to for PT) a timeout across a session. White dashed horizontal line: first Training day earning 10 rewards. Horizontal white line: acquisition threshold, when a mouse began to earn ∼20 rewards consistently in Training. Black horizontal lines: divisions between training phases. G) Heatmap of BLA ACh signaling during incorrect nose poke averaged across mice. Signal aligned as in C-F) with a selection of data from key days in the behavioral paradigm shown. From bottom to top: PT Day 1, PT Day 5, Training Day 1, Training Day 3, First Training day earning 10 rewards (white dashed horizontal line), Training Day 13, Training Day 15, Acquisition day (white horizontal line), Last Training Day, Last Extinction Day. Black horizontal lines: divisions between training phases.

**Fig. S2.3.**
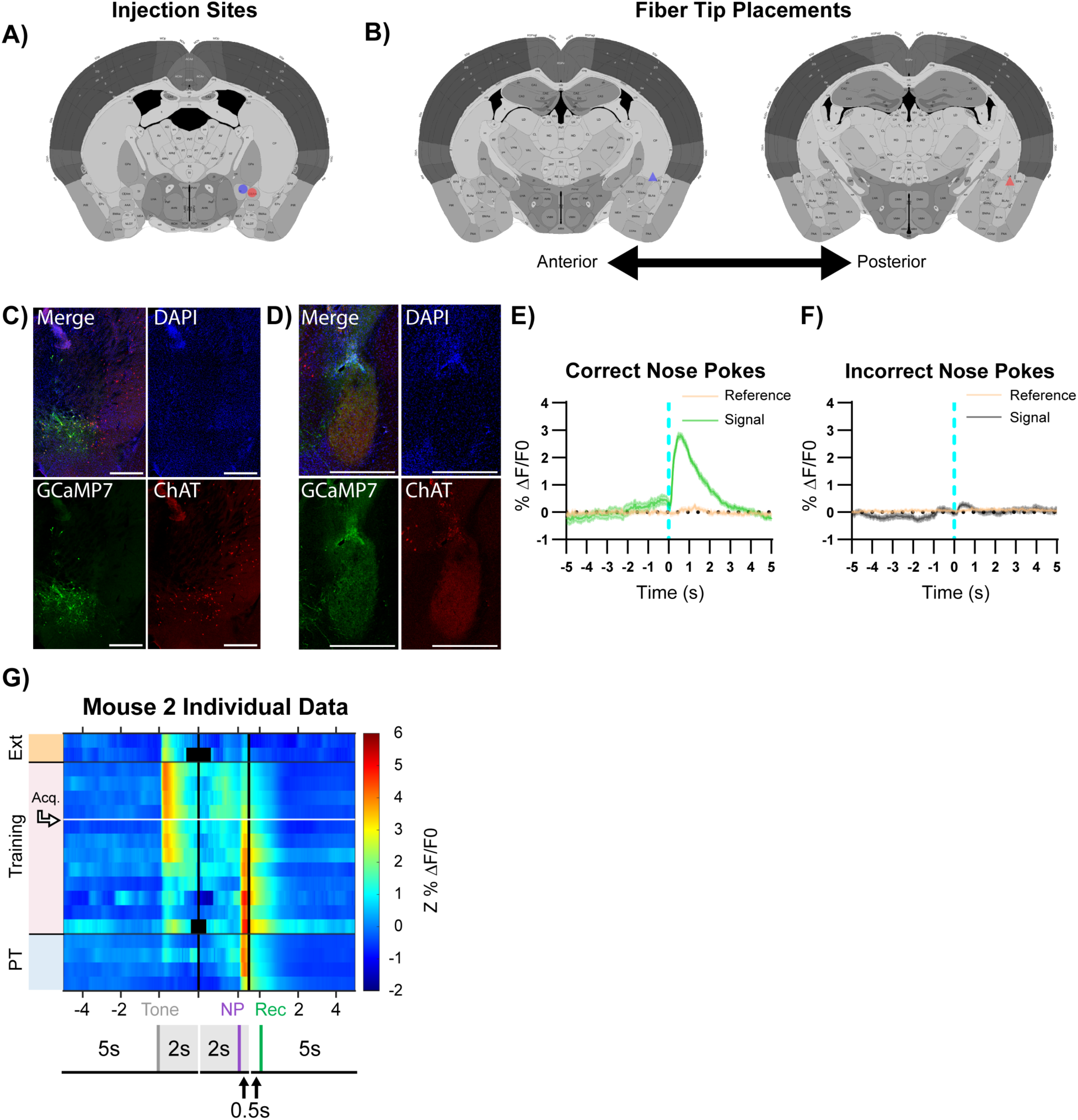
Supplemental Data for Fig. 2 F-J A) Circles indicate NBM DIO-GCaMP7s injection sites for individual mice, 1 (red), 2 (blue). B) Triangles indicate estimated optical fiber tips based on adjacent slices for individual mice. 1 (red), 2 (blue). C) Representative injection site coronal slice from Fig. 2F with channels separated. Scale = 500 µm. D) Representative fiber tip site coronal slice from Fig. 2F with channels separated. Scale = 500 µm. E) Increase in fluorescence (%ΔF/F0) following correct nose pokes is specific to the signal (465 nm, green) channel and is not observed in the reference channel (405 nm, tan). Data from Mouse 1 PT Day 4 as in Fig. 2H. Mean ± SEM, n = 42. F) Minimal increase in fluorescence (%ΔF/F0) following incorrect nose pokes. Signal (465 nm, grey) channel, reference channel (405 nm, tan). Data from Mouse 1 PT Day 4 as in Fig. 2H. Mean ± SEM, n = 101. G) Individual data for mouse 2 as shown in Fig. 2I. White horizontal line: acquisition threshold.

**Fig. S2.4.**
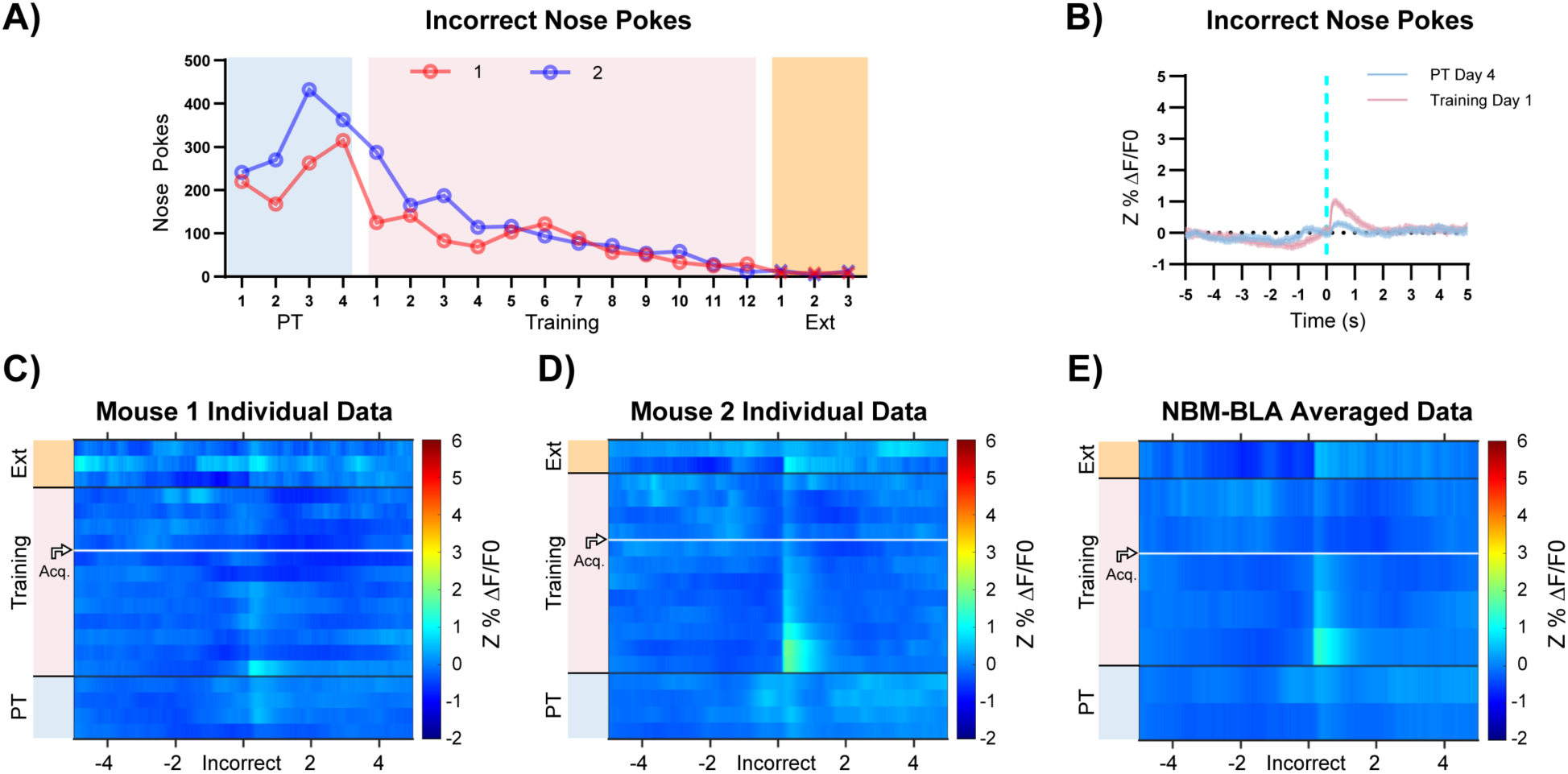
Supplemental Data for Fig. 2F-J A) Incorrect nose poking of individual mice throughout training. B) Incorrect nose pokes that yield timeouts (Training Day 1, pink line, n = 105) result in increase in NBM-BLA terminal fiber activity but incorrect nose pokes before timeouts are introduced (PT Day 4, blue line, n = 101) do not. Data from Mouse 1 as in Fig. 2H, Mean ± SEM. C-D) Individual mouse heatmaps of NBM-BLA terminal fiber activity across all training phases, aligned to incorrect nose poke. Each row is the average of incorrect nose pokes that led to (or would have led to for PT) a timeout across a session. Horizontal white line: acquisition threshold, when a mouse began to earn ∼20 rewards consistently in Training. Black horizontal lines: divisions between training phases. E) Heatmap of NBM-BLA terminal fiber activity during incorrect nose poke averaged across mice. Signal aligned as in C-D) with a selection of data from key days in the behavioral paradigm shown. From bottom to top: PT Day 1, PT Day 4, Training Day 1, Training Day 3, Training Day 6, Acquisition day (white horizontal line), Last Training Day, Last Extinction Day. Black horizontal lines: divisions between training phases.

**Fig. S2.5.**
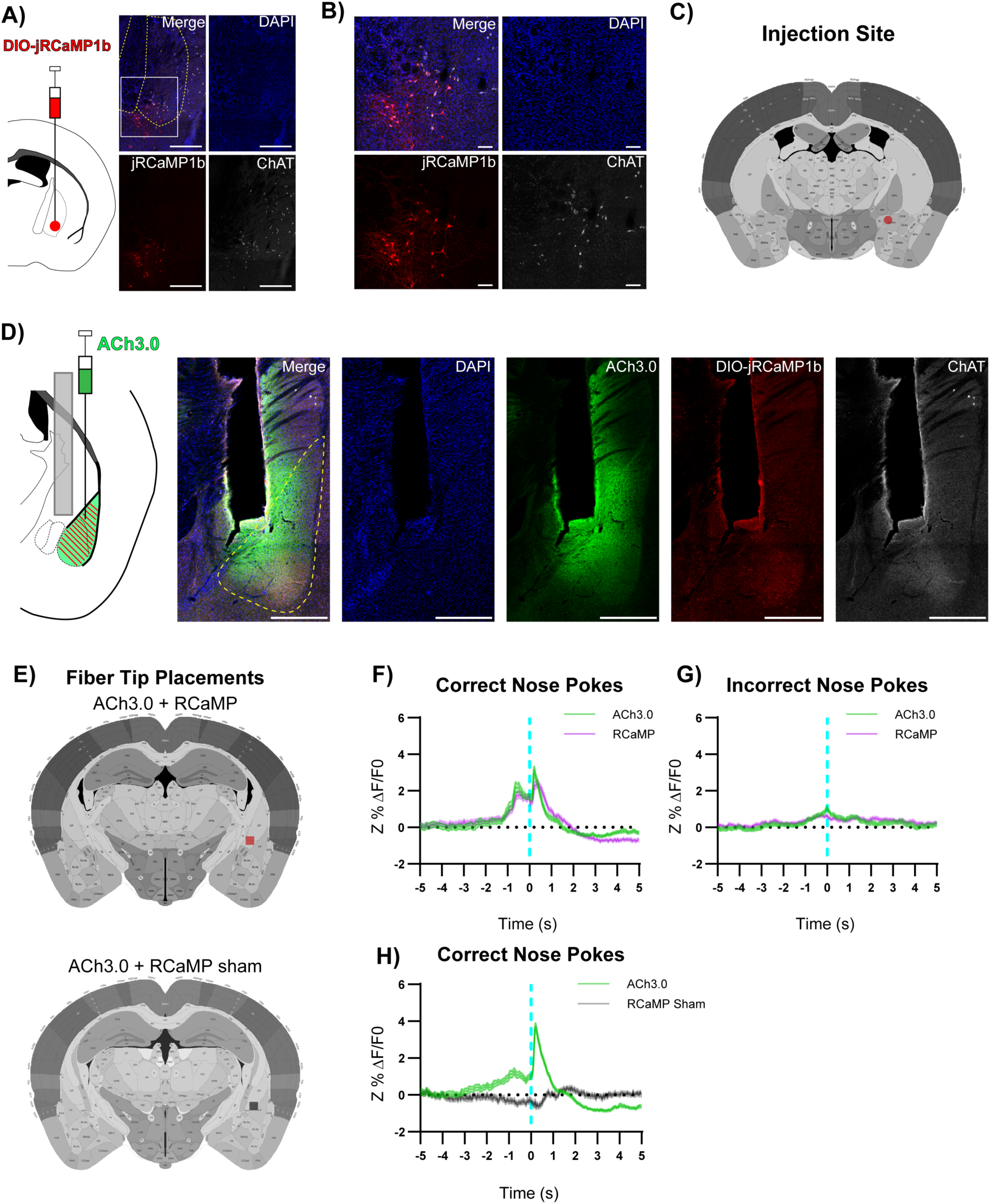
Supplemental Data for Fig. 2 A) Left: DIO-jRCaMP1b was injected in the NBM of ChAT-IRES-Cre mice. Representative coronal brain slice showing jRCaMP1b expression. Yellow dashed lines: internal capsule and globus pallidus outlines. Scale = 500 µm. White box: higher magnification area shown in B). B) Higher magnification of injection site. Scale = 100 µm. C) Circle indicates NBM DIO-jRCaMP1b injection site for mouse 1. D) ACh3.0 was injected into the ipsilateral BLA and an optical fiber was implanted above the BLA. White dashed line: BLA outline. Scale = 500 µm. E) Squares indicate optical fiber tips for individual mice. ACh3.0 + RCaMP (red), ACh3.0 + RCaMP sham (grey), F) A substantial increase in both fluorescence representing BLA ACh release (green line) and NBM-BLA cholinergic terminal activity (magenta line) coincided with correct nose pokes on last day of PT. Mean ± SEM, (n = 42). G) Minimal increase in fluorescence in either channel following incorrect nose pokes on last day of PT. Mean ± SEM, (n = 94) H) <u>jRCaMP1b signal is not simply crosstalk from ACh3.0 channel.</u> A substantial increase in fluorescence representing BLA ACh release (green line) following correct nose pokes did not necessitate signal in RCaMP sham red channel (grey line). Last day of PT. Mean ± SEM, n = 44.

**Fig. S3.1.**
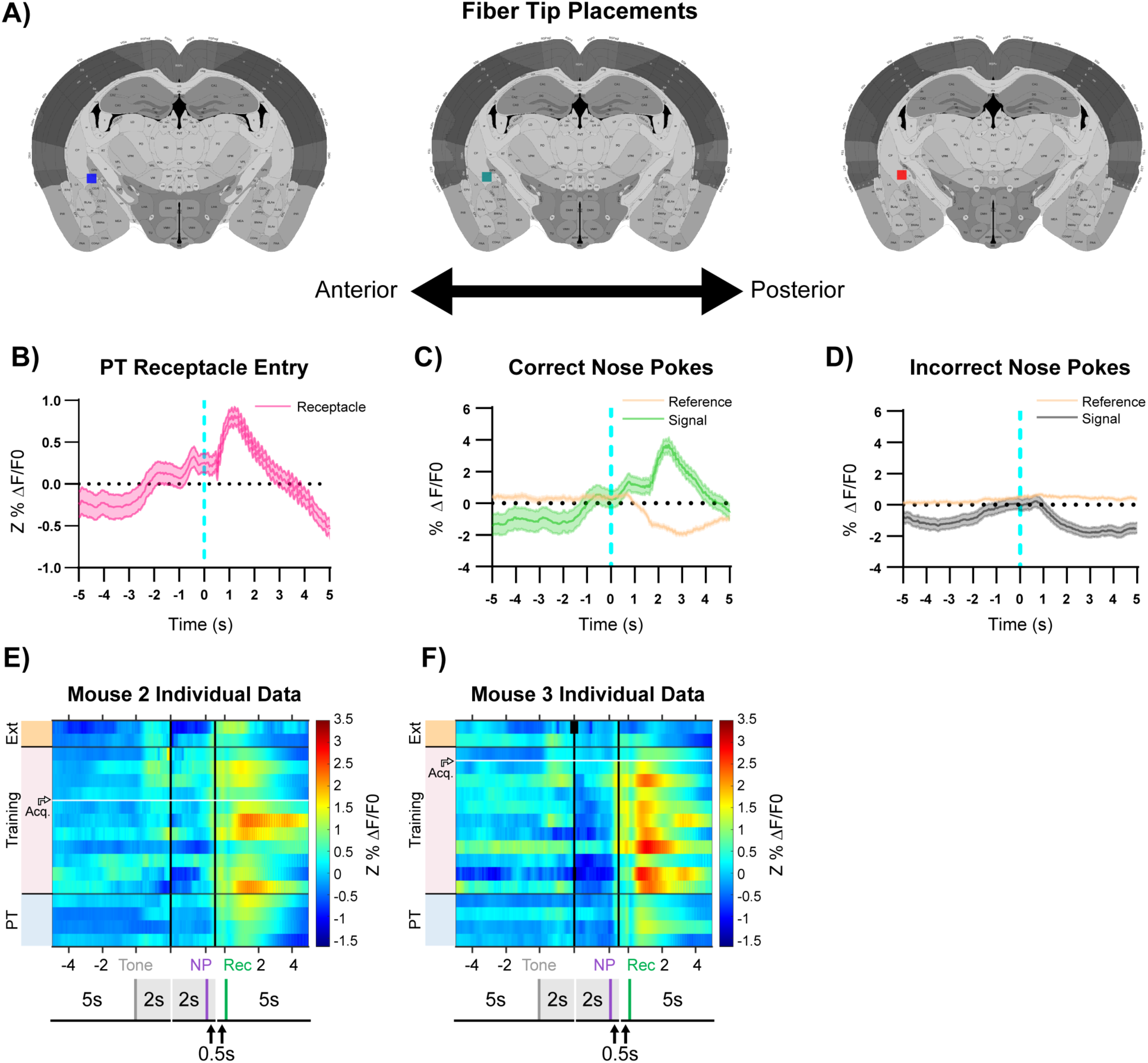
Supplemental Data for Fig. 3 A) Squares indicate optical fiber tips for individual mice. 1 (red), 2 (blue), 3 (teal). B) Increase in fluorescence (Z%ΔF/F0) during last day of PT (data shown for Mouse 1) aligns more closely to receptacle entry (reward retrieval) on rewarded trials. Mean ± SEM, n = 44. C) Increase in fluorescence (%ΔF/F0) following correct nose pokes is specific to the signal (465 nm, green) channel and is not observed in the reference channel (405 nm, tan). Dip in reference channel following correct nose poke is likely due to not acquiring at the “true” isosbestic point of GCaMP (Barnett et al., 2017; C. K. Kim et al., 2016; Sych et al., 2019)). Data from Mouse 1, PT Day 4 as in Fig. 3B. Mean ± SEM, n = 44. D) Decrease in fluorescence (%ΔF/F0) following incorrect nose pokes is seen in signal channel (465 nm, grey), but not reference channel (405 nm, tan). Data from Mouse 1, PT Day 4 as in Fig. 3B. Mean ± SEM, n = 141. E-F) Individual data for mice not shown in Fig. 3D. White horizontal line: acquisition threshold.

**Fig. S3.2.**
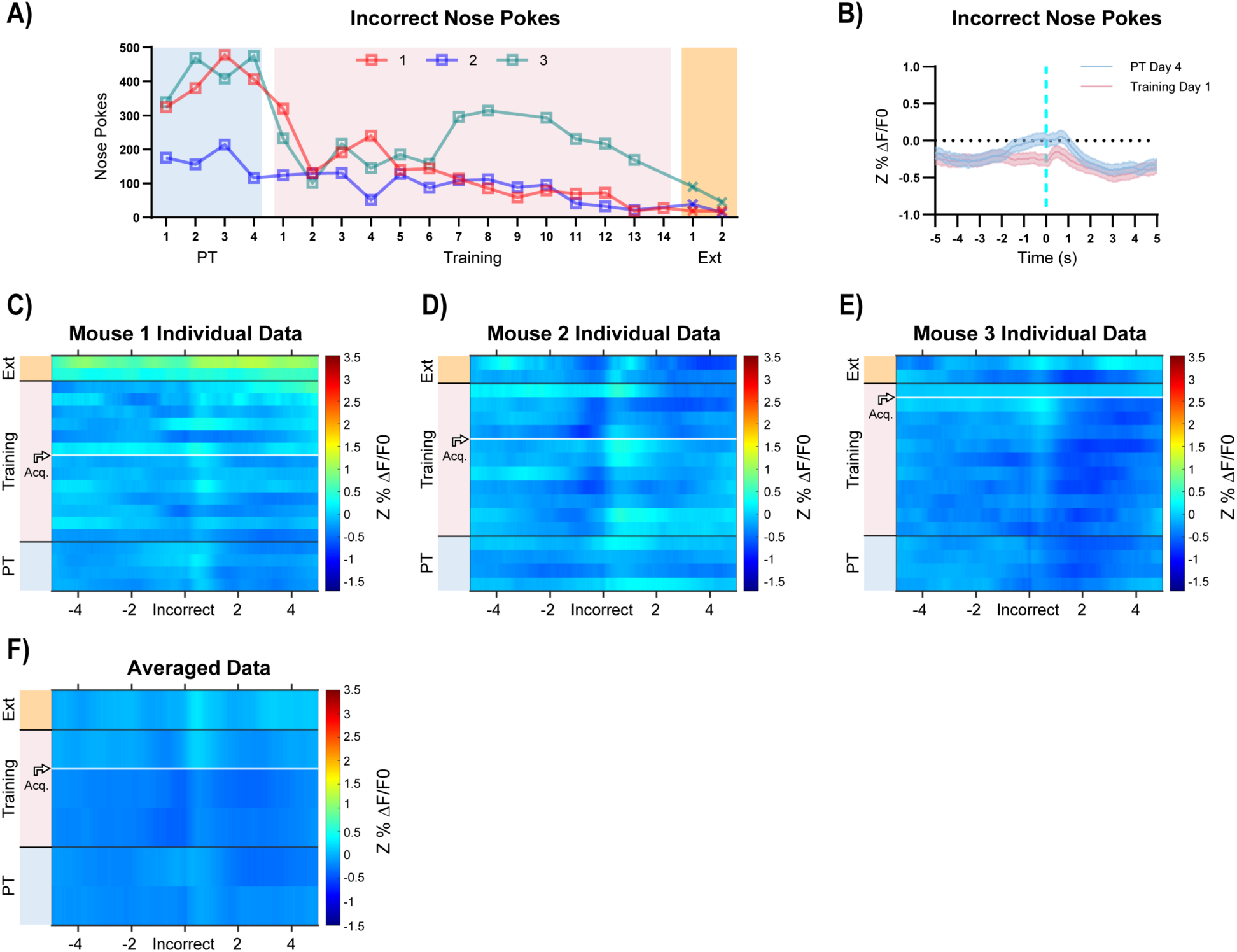
Supplemental Data for Fig. 3 A) Incorrect nose pokes of individual mice throughout training. B) Both incorrect nose pokes that yield timeouts (Training Day 1, pink line, n = 124) and incorrect nose pokes before timeouts are introduced (PT Day 4, blue line, n = 141) result in a decrease in BLA principal neuron activity. Data from Mouse 1 as in Fig. 3B, Mean ± SEM. C-E) Individual mouse heatmaps of BLA principal neuron activity across all training phases, aligned to incorrect nose poke. Each row is the average of incorrect nose pokes that led to (or would have led to for PT) a timeout across a session. Horizontal white line: acquisition threshold, when a mouse began to earn ∼20 rewards consistently in Training. Black horizontal lines: divisions between training phases. F) Heatmap of BLA principal neuron activity during incorrect nose poke averaged across mice. Signal aligned as in C-E) with a selection of data from key days in the behavioral paradigm shown. From bottom to top: PT Day 1, PT Day 4, Training Day 1, Training Day 3, Acquisition day (white horizontal line), Last Extinction Day. Black horizontal lines: divisions between training phases.

**Fig. S4.1.**
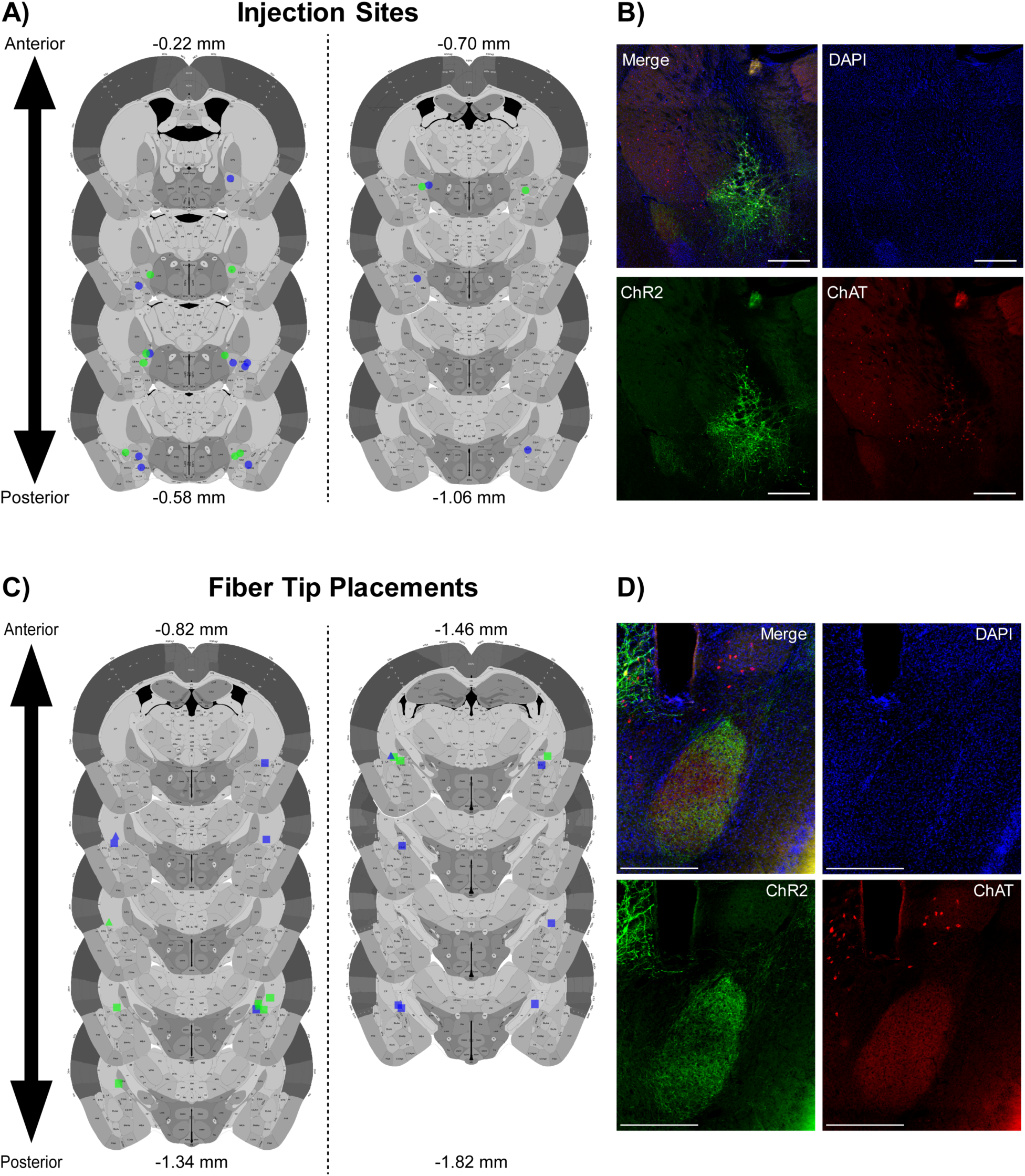
Injection Sites Optical Fiber Placements for Fig. 4 A) Circles indicate NBM injection sites for individual mice, EYFP (green) and ChR2 (blue). Anterior/Posterior position relative to Bregma indicated. B) Representative injection site coronal slice from Fig. 4A with channels separated. Scale = 500 µm. C) Squares indicate observable optical fiber tips for individual mice, EYFP- (green) and ChR2- expressing mice (blue). Triangles indicate estimated optical fiber tips based on adjacent slices. Anterior/Posterior position relative to Bregma indicated. D) Representative fiber tip site coronal slice from Fig. 4A with channels separated. Scale = 500

**Fig. S4.2.**
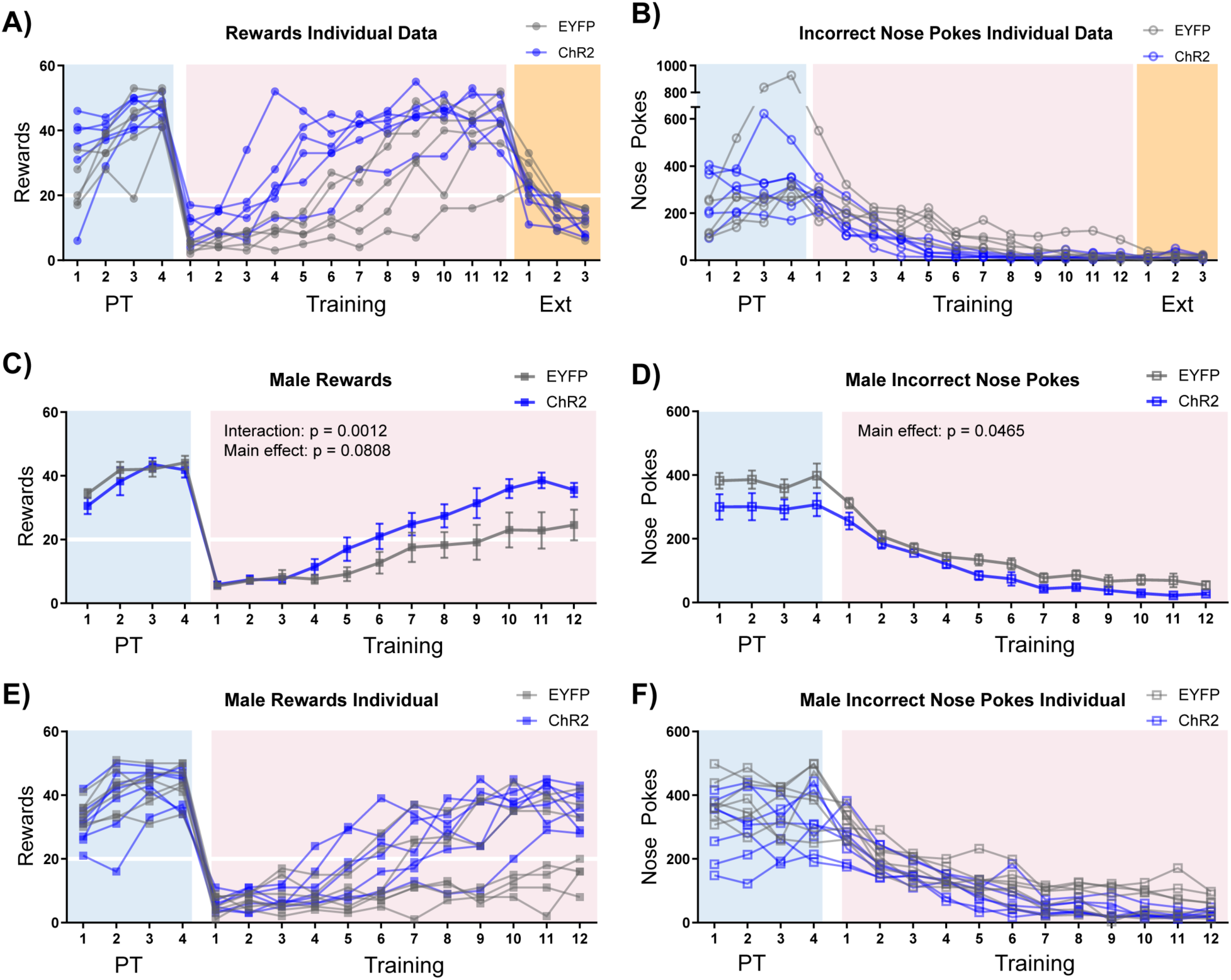
Individual Data for Fig. 4 and Males A) Rewards earned for individual mice from Fig. 4E. Horizontal white line: acquisition threshold, when a mouse began to earn ∼20 rewards consistently in Training. B) Incorrect nose pokes for individual mice from Fig. 4F. C) Optical stimulation of ChAT^+^ NBM-BLA terminal fibers (ChR2-expressing mice, blue squares) had a similar effect on rewards earned during Training in male mice compared to female mice. Mean ± SEM, EYFP: n = 7, ChR2: n = 7. Horizontal white line: acquisition threshold, when a mouse began to earn ∼20 rewards consistently in Training. D) Optical stimulation of ChAT^+^ NBM-BLA terminal fibers (ChR2-expressing mice, blue squares) had a similar effect on incorrect nose pokes during Training in male mice compared to female mice. Mean ± SEM, EYFP: n = 7, ChR2: n = 7. E) Individual data for graph shown in C). F) Individual data for graph shown in D).

**Fig. S4.3.**
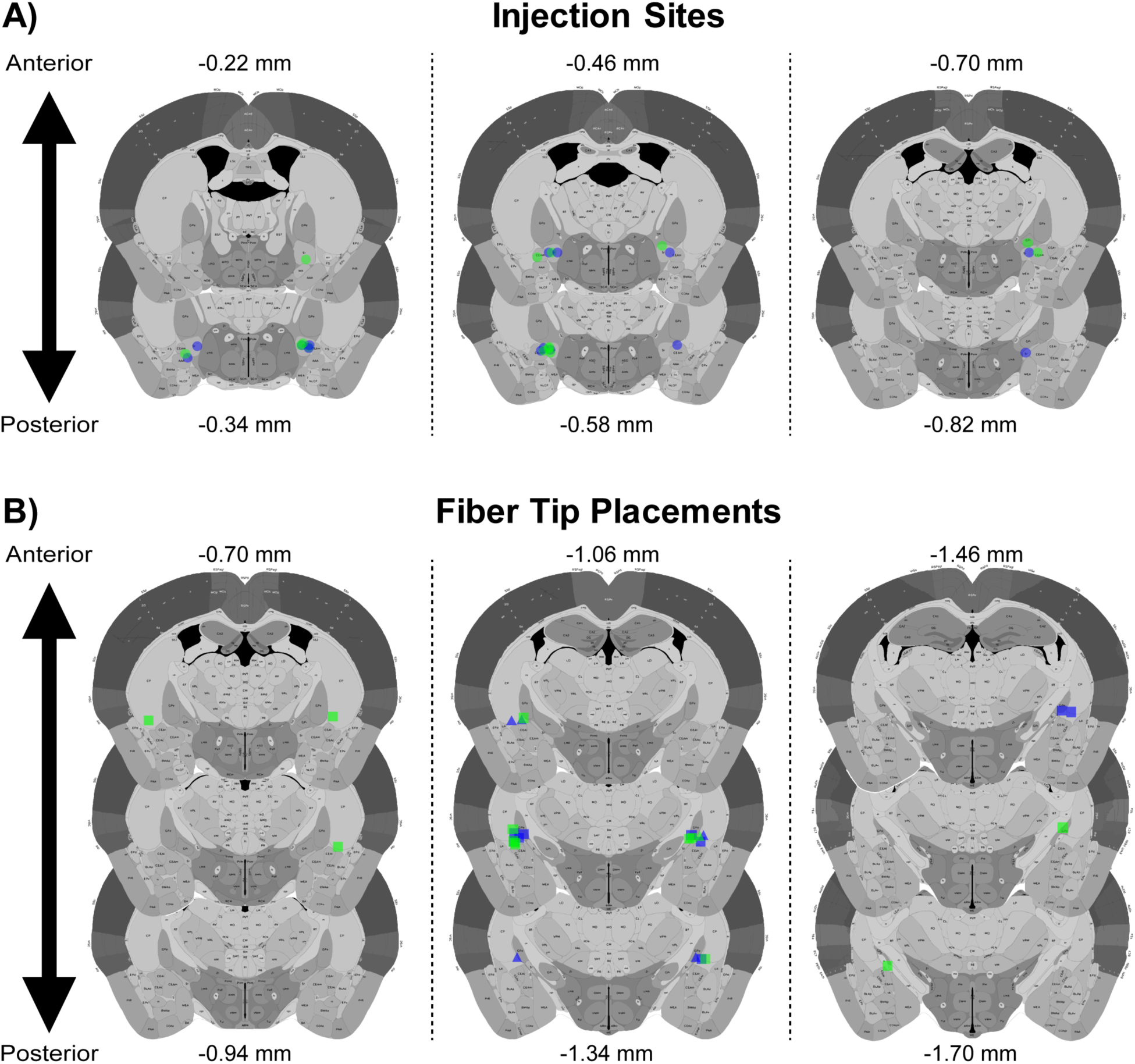
Injection Sites and Optical Fiber Placements for Fig. S4.2C-F A) Circles indicate NBM injection sites for individual mice, EYFP- (green) and ChR2-expressing mice (blue). Anterior/Posterior position relative to Bregma indicated. B) Squares indicate observable optical fiber tips for individual mice, EYFP- (green) and ChR2- expressing mice (blue). Triangles indicate estimated site of optical fiber tips based on adjacent slices. Anterior/Posterior position relative to Bregma indicated.

**Fig. S4.4.**
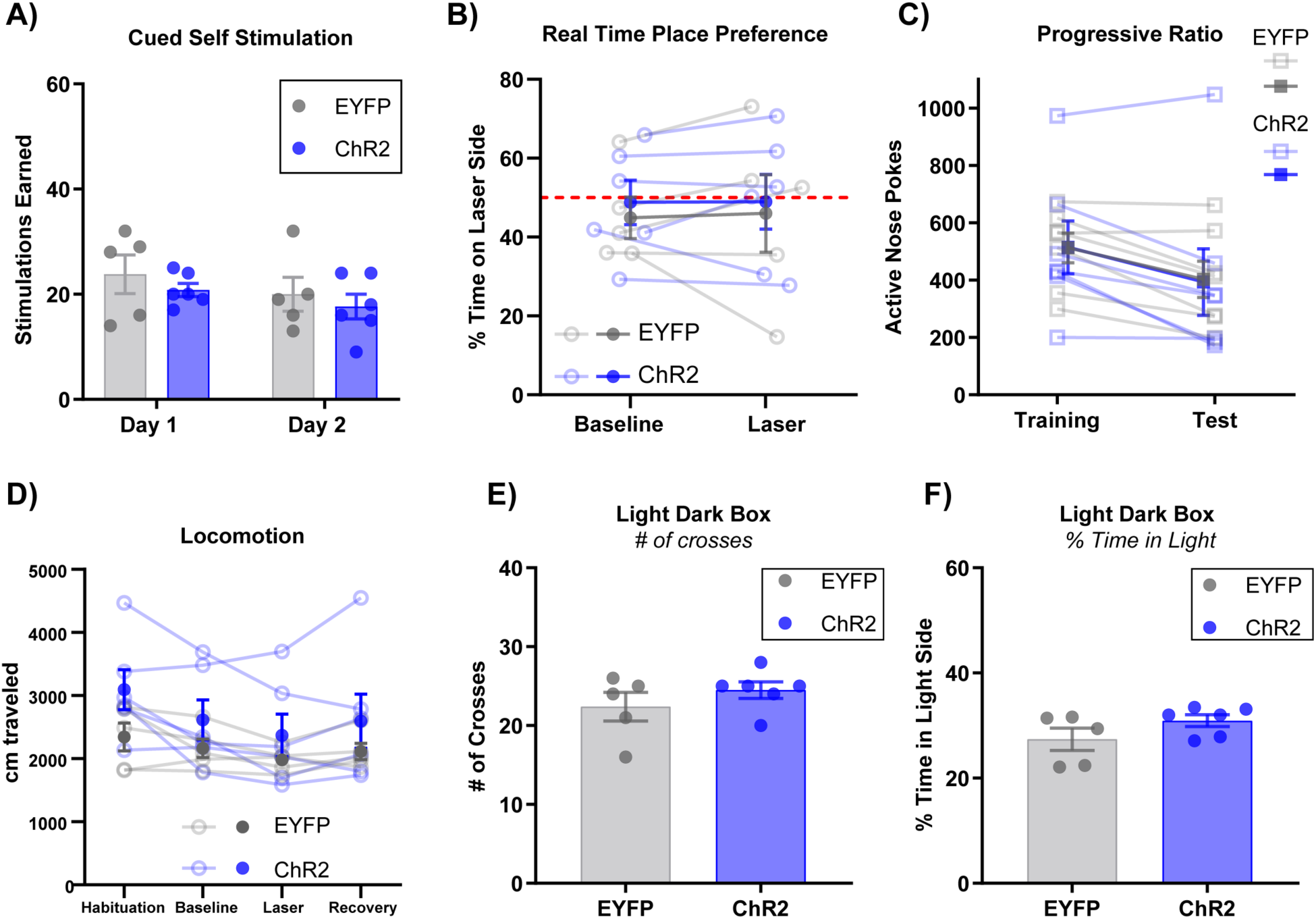
Additional Behavioral Assays for Fig. 4 A) Stimulation of ChAT^+^ NBM-BLA terminal fibers did not support self-stimulation. Mice were allowed to nose poke for 2 sec of stimulation in the Training paradigm. Data for female mice from Fig. 4 + **Fig. S4.1-S4.2A-B.** B) Stimulation of ChAT^+^ NBM-BLA terminal fibers did not support real time place preference. Mice were allowed to move freely between two sides of an empty cage with distinct floor contexts for 15 min. Data are reported as percent time spent on the laser-paired side. Closed circles: Mean ± SEM, open circles: data for individual mice. Data for female mice from Fig. 4 + **Fig. S4.1-S4.2 A-B.** C) Stimulation of ChAT^+^ NBM-BLA terminal fibers during a progressive ratio test did not affect active nose poking. Closed squares: Mean ± SEM, open squares: individual mice. Data for male mice from **Fig. S4.2C-F + S4.3**. D) There were no differences between EYFP- and ChR2-expressing mice in locomotor activity. X-axis ticks = 5 min bins, Laser = 5 min of 20 sec on/off optical stimulation. Closed circles: Mean ± SEM, open circles: data for individual mice. Data for female mice from Fig. 4 + **Fig. S4.1-S4.2 A-B.** E-F) No difference in behavior was seen between EYFP- and ChR2-expressing mice on any measures in the Light/Dark Box Test. Data for female mice from Fig. 4 + **Fig. S4.1-S4.2 A-B.**

**Fig. S5.1.**
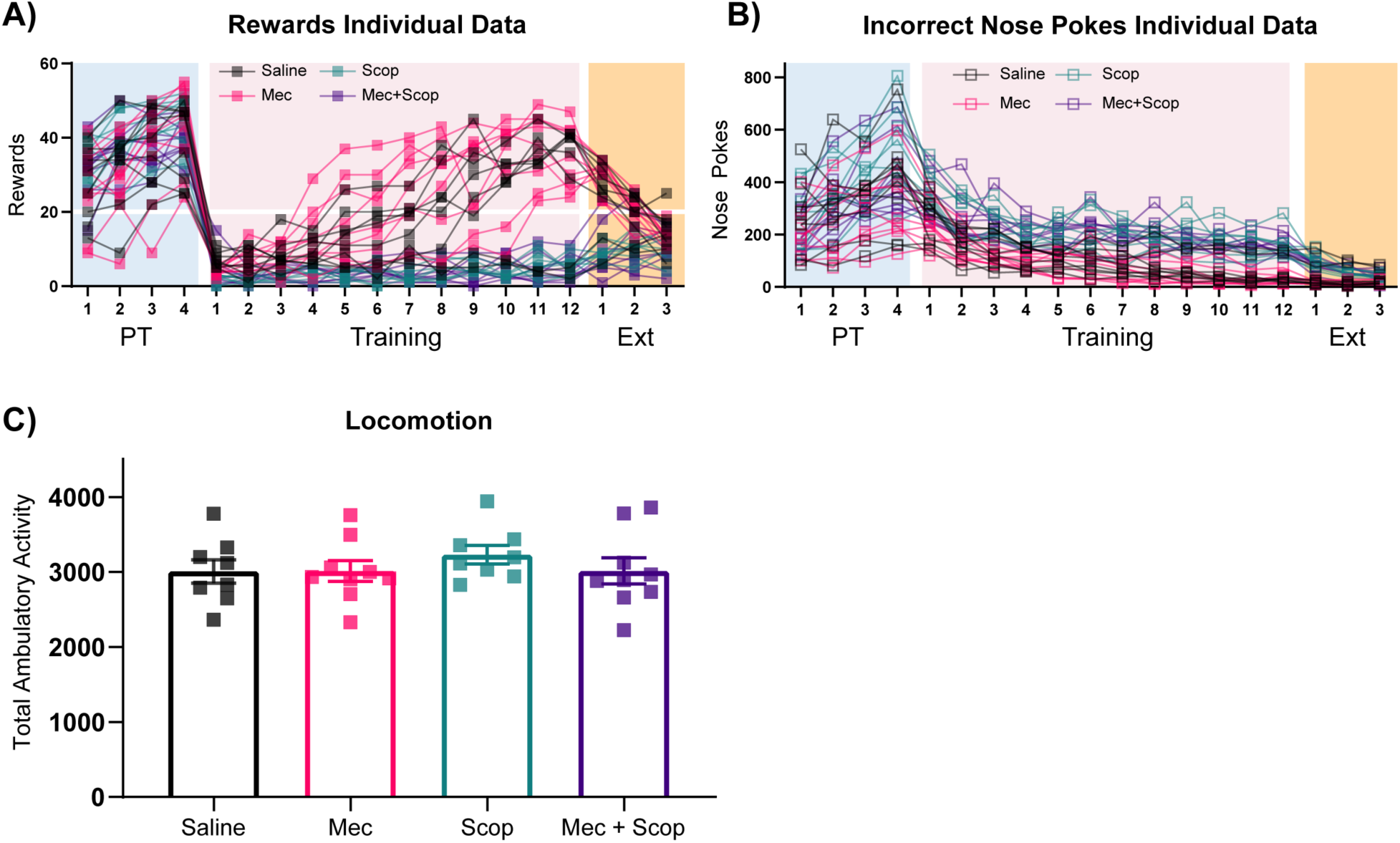
Individual Data for Fig. 5 and Locomotion A) Rewards earned for individual mice from Fig. 5B. Horizontal white line: acquisition threshold, when a mouse began to earn ∼20 rewards consistently in Training. B) Incorrect nose pokes for individual mice from Fig. 5C. C) There were no differences in locomotion for antagonists.

**Fig. S6.1.**
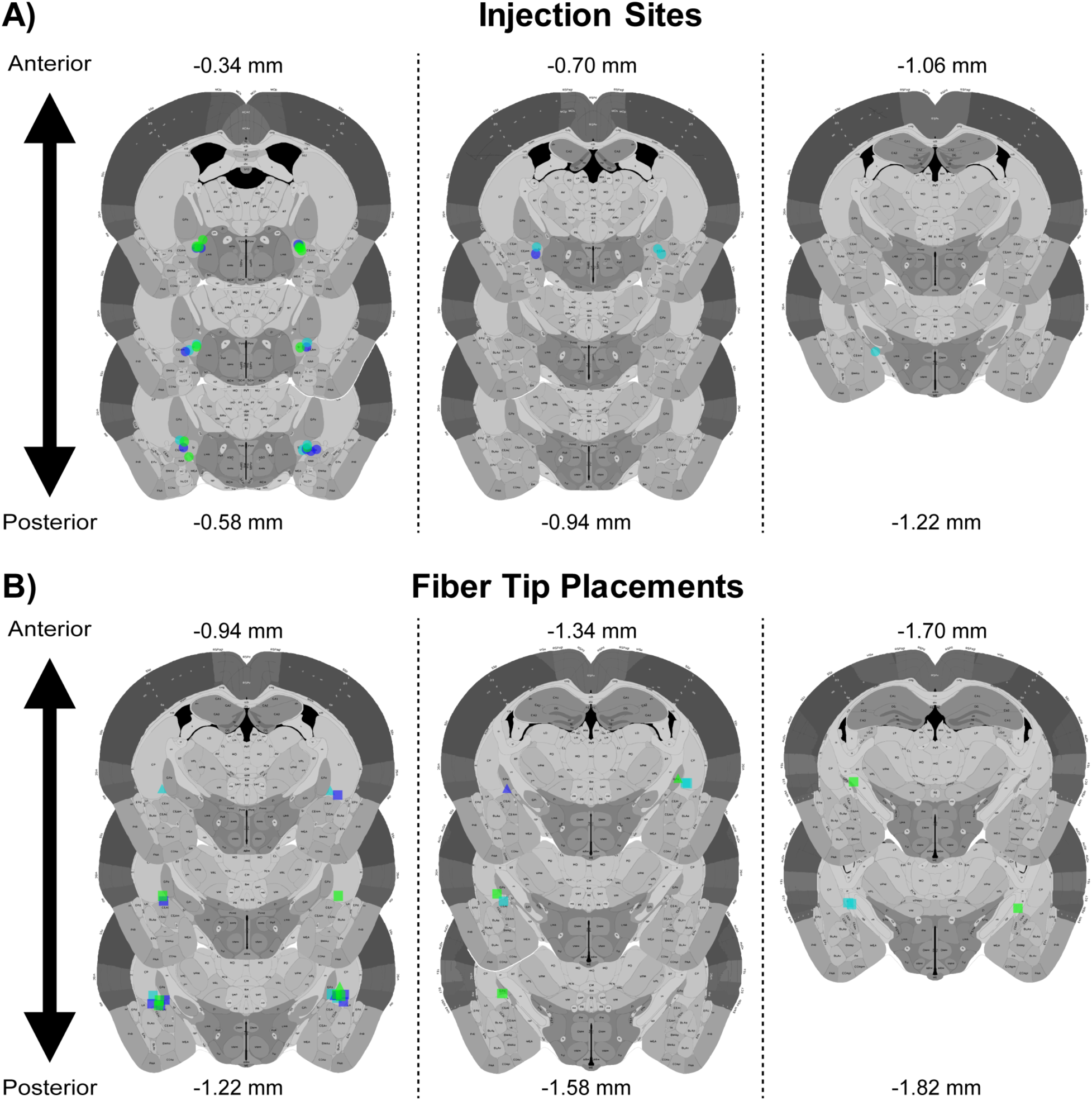
Injection Sites and Optical Fiber Placements for Fig. 6 A) Circles indicate NBM injection sites for individual mice, EYFP-expressing (green), ChR2-expressing Contingent (blue), and ChR2-expressing Yoked mice (cyan). Anterior/Posterior position relative to Bregma indicated. B) Squares indicate observable optical fiber tips for individual mice, EYFP-expressing (green), ChR2-expressing Contingent (blue), and ChR2-expressing Yoked mice (cyan). Triangles indicate estimated site of optical fiber tips based on adjacent slices. Anterior/Posterior position relative to Bregma indicated.

**Fig. S6.2.**
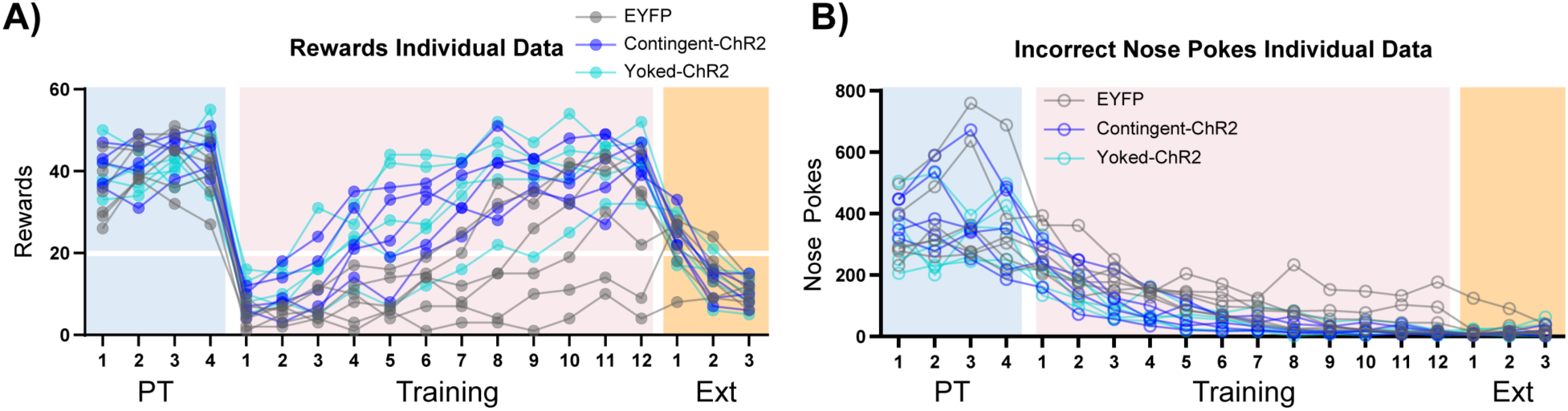
Individual Data for Fig. 6 A) Rewards earned for individual mice from Fig. 6B. Horizontal white line: acquisition threshold, when a mouse began to earn ∼20 rewards consistently in Training. B) Incorrect nose pokes for individual mice from Fig. 6C.

